# Structure Activity Relationship of the Stem Peptide in Sortase A mediated Ligation from *Staphylococcus aureus*

**DOI:** 10.1101/2022.05.17.492301

**Authors:** Alexis J. Apostolos, Joey J. Kelly, George G. Ongwae, Marcos M. Pires

## Abstract

The surfaces of most Gram-positive bacterial cells, including that of *Staphylococcus aureus* (*S. aureus*), are heavily decorated with proteins that coordinate cellular interactions with the extracellular space. In *S. aureus*, sortase A is the principal enzyme responsible for covalently anchoring proteins, which display the sorting signal LPXTG, onto the peptidoglycan (PG) matrix. Considerable efforts have been made to understand the role of this signal peptide in the sortase-mediated reaction. In contrast, much less is known about how the primary structure of the other substrate involved in the reaction (PG stem peptide) could impact sortase activity. To assess the sortase activity, a library of synthetic analogs of the stem peptide that mimic naturally existing variations found in the *S. aureus* PG primary sequence were evaluated. Using a combination of two unique assays, we showed that there is broad tolerability of substrate variations that are effectively processed by sortase A. While some of these stem peptide derivatives are naturally found in mature PG, they are not known to be present in the PG precursor, lipid II. These results suggest that sortase A could process both lipid II and mature PG as acyl-acceptor strands, which has not been previously described.

## Introduction

Many pathogenic Gram-positive organisms covalently anchor virulence proteins to the surface of their cell wall, a highly crosslinked meshwork known as peptidoglycan (PG). The PG is a thick outer layer that surrounds the cytoplasmic membrane in Gram-positive organisms (**Figure 1A**). PG is typically comprised of the disaccharide *N*-acetylglucosamine (GlcNAc) and *N*-acetylmuramic acid (MurNAc) as a monomermic unit that is connected to form a polymeric scaffold. Attached to each MurNAc unit is a unique, short peptide (called the stem peptide) composed of up to five amino acids in length. The canonical pentapeptide sequence is typically L-Ala-*iso*-D-Glu-L-Lys (or *meso*-diaminopimelic acid [*m*-DAP])-D-Ala-D-Ala (**Figure 1B**). It is now well established that the primary sequence can have variation within a single bacterial cell but also across different organisms, especially at the third position.^*1*^ A large fraction of bacterial species has an *m*-DAP residue at the third position^*2*^ and most of the remaining have L-Lys in this position. A common observation for organisms with L-Lys in the third position, is that additional amino acids are often connected to amino group on the L-Lys sidechain, and these additional amino acids are collectively known as cross-bridging amino acids. Cross-bridging amino acids can include – but are not limited to – L-Ala, D-Ser, D-iAsx (where Asx indicates aspartic acid or asparagine), and Gly.^*2, 3*^

**Figure 1.**
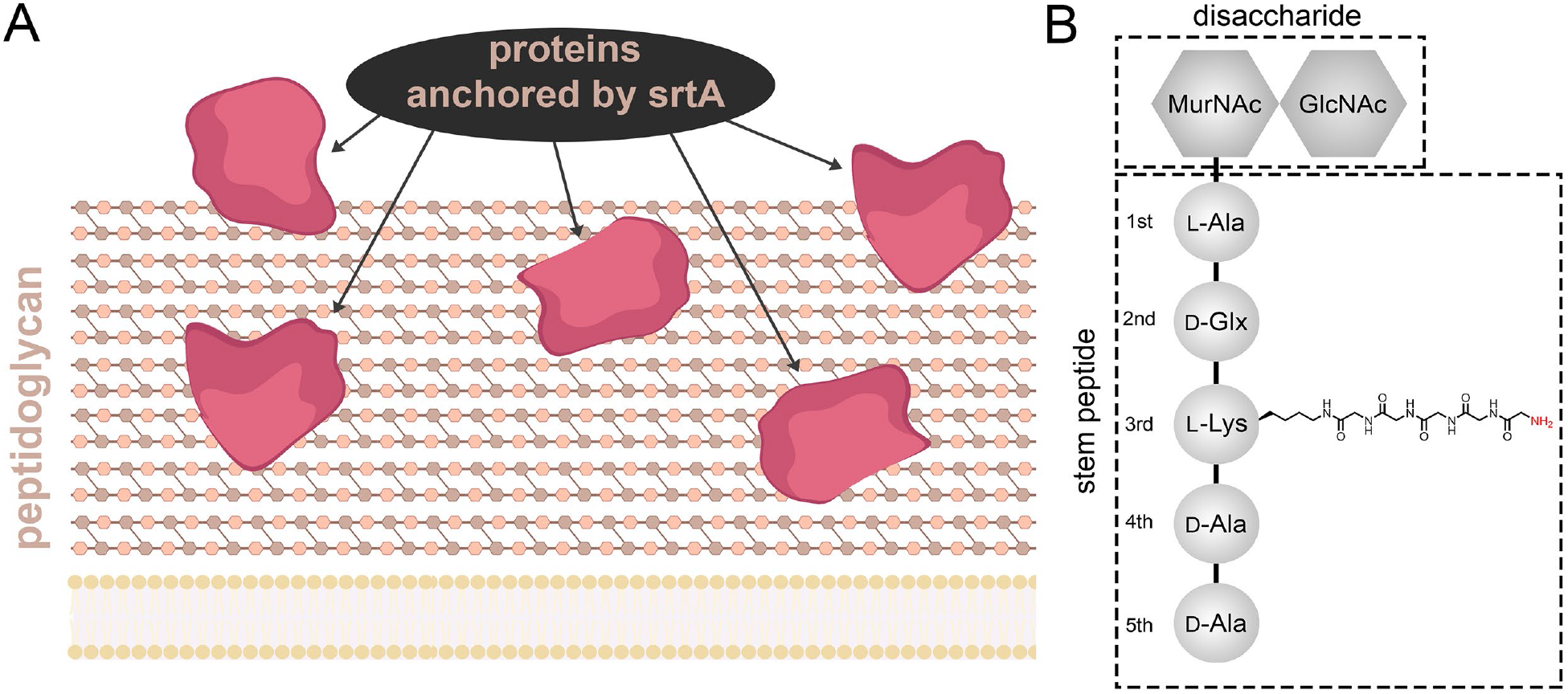
(A) Schematic representation of the surface of *S. aureus*. The enzyme srtA covalently anchors proteins onto the peptidoglycan layer, which are imbedded within and displayed onto the surface of these bacteria. (B) Canonical monomeric unit of peptidoglycan for *S. aureus*, which includes the disaccharide backbone and the stem peptide. Site of acyl-acceptor nucleophile is shown in red.

The location of the PG matrix provides an opportunity for Gram-positive organisms to stage proteins that support functions that are important for survival and proliferation. A primary mode of protein display involves the covalent anchoring of proteins within the PG matrix, a reaction carried out by sortases. *Staphylococcus aureus* (*S. aureus)* is a prominent example of a human pathogen that utilizes sortase-mediated anchoring to acquire iron, promote adhesion, and subvert host immune response.^*4*^ More specifically, immune evasion occurs by the display of protein A, which binds the Fc region of immunoglobin G (IgG) during infection to reduce recognition by the host immune system.^*5-7*^ Sortase enzymes are transpeptidases that recognize a specific sorting sequence present in the protein for eventual anchoring within the PG.^*8, 9*^ In *S. aureus*, the sorting sequence for processing by sortase A (srtA) is LPXTG, whereby X is any amino acid.^*10*^ The cysteine residue in the active site of srtA clips the peptide bond between T and G of the sorting sequence, thus forming a thioacyl intermediate between sortase and the threonine of the LPX<u>T</u>-containing protein.^*11, 12*^ This thio-acyl intermediate undergoes a nucleophilic attack by the *N*-terminal amino group, which is found on the pentaglycine (G_5_) cross-bridge of *S. aureus* PG (**Figure 2**).^*13*^ A new amide bond is created between the protein and the PG pentaglycine cross-bridge, allowing for covalent display of virulent proteins on the bacterial cell surface.^*14, 15*^

**Figure 2.**
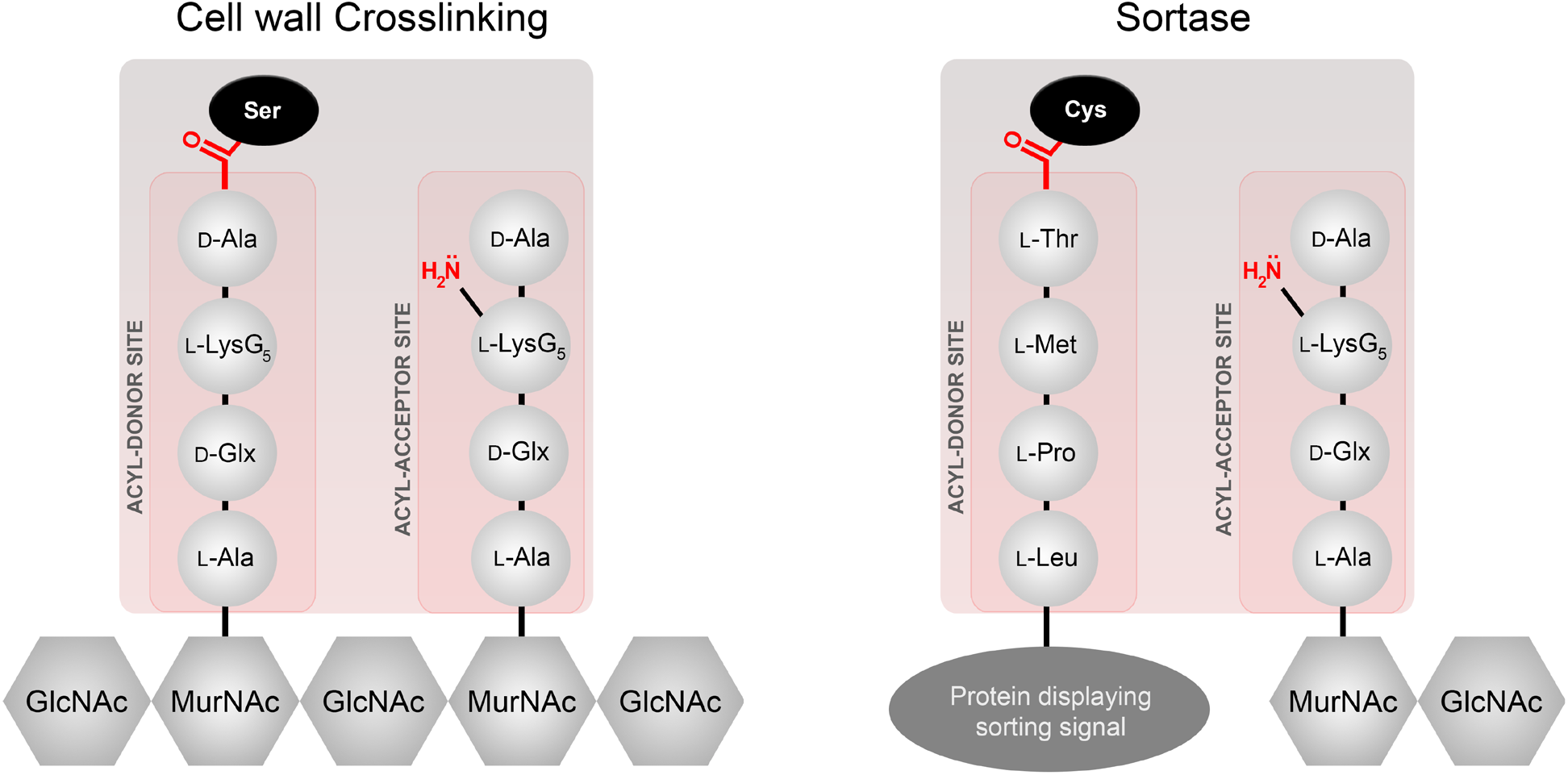
Similarity in the two reactions that use a common acyl-acceptor strand. In *S. aureus*, cell wall crosslinking of neighboring strands is performed by PBPs and Ldts (left). A similar reaction pathway for the second step of the reaction is observed for srtA, with the primary difference being the acyl-donor strand.

Due to the fact that srtA aids *S. aureus* in host-colonization and altered host-immune response, it is considered a promising anti-virulence target. The deletion of srtA in *S. aureus* disrupts the establishment of infection in mouse models.^*16*^ These results have spurred considerable efforts to discover effective srtA inhibitors.^*17-19*^ Additionally, the transpeptidase activity of sortase has been harnessed for use as a general ligation tool outside the context of bacterial cell surfaces. This has been demonstrated by extensive work whereby various synthetic nucleophiles have been used, as well as a multitude of LPTXG-containing scaffolds, in place of the natural substrates found within the bacterial cell, to ligate molecules of interest.^*20-24*^ Interestingly, the transpeptidase reaction performed by sortase has functional similarity to the transpeptidases responsible for PG crosslinking, carried out by either penicillin binding proteins (PBPs) or L,D-transpeptidases (Ldts). These transpeptidase enzymes use two neighboring strands of PG to generate 4-3 (between the 4^th^ position of the donor strand and 3^rd^ position of the acceptor strand) or 3-3 (between the 3^rd^ position on both strands) crosslinks.^*25, 26*^ The primary difference between these two types of transpeptidases occurs in the first half of the reaction. Whereas PG crosslinking enzymes process a stem peptide as the acyl-donor strand, srtA will instead process the LPXTG (**Figure 2**). The second half of the reaction – the nucleophilic attack from amino group on the G_5_ crossbridge – potentially utilizes the same substrate.

Given the essentiality of PG crosslinking, as demonstrated by the number and diversity of small molecule antibiotics that disrupt this pathway, it is possible that dysregulated processing of the same substrate (acyl-acceptor strand) could lead to loss of cell wall integrity. More specifically, we wondered whether the regulation of acyl-acceptor utilization could be inherently dictated by the primary sequence of the stem peptide. It is well established that the stem peptide in *S. aureus*, and for most bacteria whose PG have been analyzed, can vary in length (penta-, tetra-, and tri-peptides all retain the acyl-accepting crossbridge)^*27, 28*^ and amidation states of D-iGlu.^*29, 30*^ To investigate this, we systematically measured sortase activity of srtA *in vitro* with variations of possible acyl-acceptor stem peptides endogenously found within the PG of *S. aureus*. Interestingly, there was not a strong preference within the stem peptide primary sequence variations. It had been described that the acyl-donor strand for srtA is lipid II.^*14*^ Our results suggest that it may be possible that the acyl-donor strand comes from a more mature PG scaffold. This could indicate that sortase-mediated transpeptidase activity can be decoupled from lipid II pools.

## Results & Discussion

We first set out to measure the activity of sortase A using gel fluorescence analysis.^*31-33*^ We synthesized (using standard solid phase peptide coupling procedures) a truncated substrate of the acyl-acceptor strand GGGK(5,6-carboxytetramethylrhodamine, TAMRA; GGGK(tmr))^*34-36*^ whereby a red-fluorophore is connected to the sidechain of lysine. Additionally, we expressed and purified a green fluorescent protein (GFP) containing a *C*-terminal LPETG sorting signal to serve as the acyl-donor strand. Reactions were carried out using standard conditions and following this incubation period, the reaction contents were boiled in sodium dodecyl sulfate (SDS) to denature the GFP. Accordingly, the fluorescence observed is expected to come solely from a productive transpeptidation of the TAMRA-containing strand onto the larger molecular weight of the modified GFP. An expected product band of ∼29 kDa was anticipated for GFP-LPETGGGK(tmr) products and was observed by gel analysis (**Figure 3A**). Reactions were also set up in the absence of each reaction constituent (e.g., no srtA, no GFP, or no GGGK(tmr)) as negative controls. Using this set up, a time point assay was also performed, where the reaction with all elements was quenched at 1, 2, 4, 8 h and displayed an increase in fluorescence as time progressed (**Figure S1**).

**Figure 3.**
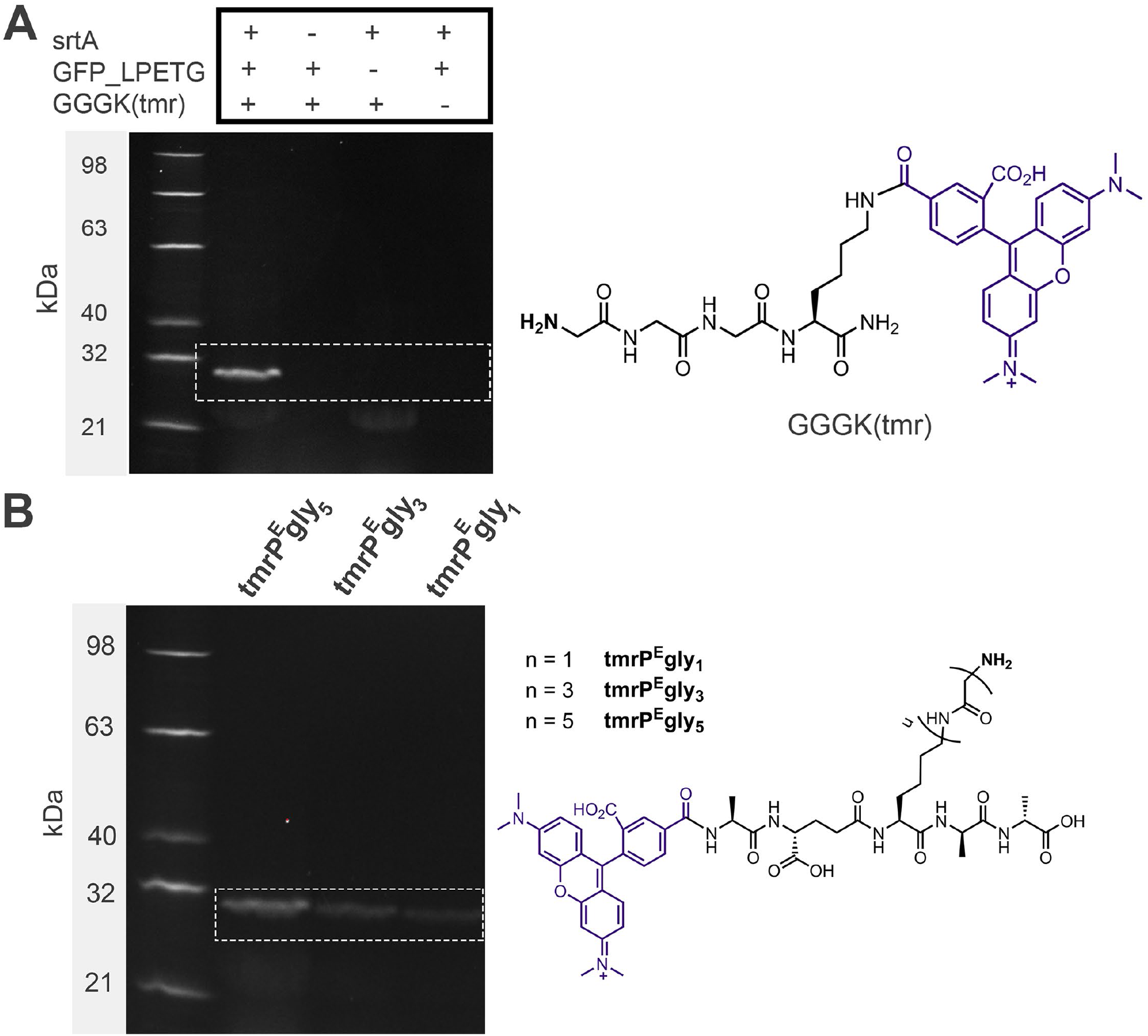
(A) Gel fluorescence analysis of GGK(tmr) in the various stated conditions. Reaction was performed at 37 oC and allowed to reaction for 8h, then loaded on an SDS-PAGE. (B) Gel fluorescence analysis of GFP_LPETG and srtA in the presence of **tmrP**^**E**^**gly**_**1**_, **tmrP**^**E**^**gly**_**3**_, or **tmrP**^**E**^**gly**_**5**_. Reaction was performed at 37 °C and allowed to reaction for 2 h, then loaded on an SDS-PAGE.

With reaction parameters optimized, we assembled a panel of stem peptide mimetic probes that would allow us to interrogate the necessity of naturally occurring peptide lengths or modifications for srtA activity. Our group, and others, previously showed that synthetic acyl-acceptor strands modified with an *N*-terminal fluorophore are well tolerated by cell wall crosslinking enzymes (PBPs and Ldts).^*37-43*^ We envisioned that a similar tolerability would enable the structure-activity relationship of srtA. As wildtype *S. aureus* displays pentaglycine cross-bridge (G_5_) appended to its 3^rd^ position lysine sidechain, we first sought to determine the importance of the glycine cross-bridge length. Collectively, biosynthesis of this cross-bridge is performed by enzymes in the *fem* gene family (*femA, femB, femAB, femAX*).^*3, 44-46*^ It had been shown that non-pentaglycyl cross-bridges have the ability to ligate surface proteins to the PG, although the natural nucleophilic cross-bridge is understood to be preferred.^*13, 47*^ We synthesized a penta-stem peptide with a TAMRA-modified *N*-terminus and varied the glycine bridge length as 5 (**tmrP**^**E**^**gly**_**5**_), 3 (**tmrP**^**E**^**gly**_**3**_), or 1 (**tmrP**^**E**^**gly**_**1**_) glycine(s) (**Figure 3**). Reactions were set up with each of the probes, srtA, and GFP-LPETG and analyzed by in-gel fluorescence. Interestingly, there appeared to be a slight increase in transpeptidation with the native glycine cross-bridge at 2 h. A distinguishably brighter signal for **tmrP**^**E**^**gly**_**5**_ was observed over both **tmrP**^**E**^**gly**_**3**_ and **tmrP**^**E**^**gly**_**1**_ (**Figure 3B**). The differences in fluorescence intensity became smaller with longer incubation time, which may indicate that subtle differences in srtA activity may only be observable in early time points prior to reaching reaction equilibrium (**Figure S3**). Further, we anticipated that this reaction product could be distinguished by a shift in mass from the GFP-LPETG alone (∼ 29 kDa) and the mass that corresponded with the newly ligated product (∼29 kDa + 1.2 kDa). As examined by MALDI-TOF, this mass shift was distinguishable, highlighting that the sortase reaction successfully ligated **tmrP**^**E**^**gly**_**5**_ onto GFP-LPETG (**Figure S4**).

While the in-gel fluorescence method could qualitatively discern large changes in transpeptidation preferences, we realized that the structure-activity relationship in the acyl-acceptor chain may be subtle. Therefore, it was important to develop an assay with a higher level of signal-to-noise ratio and sensitivity. We reasoned that it is possible load the sorting signal onto a flow cytometry-compatible bead and co-incubate with srtA and fluorescent stem peptide analogs linked to a fluorophore for analysis by flow cytometry (**Figure 4**). Upon transpeptidation of sortase, the fluorescent handle would be covalently attached to the bead, which could be readily and efficiently analyzed by flow cytometry. The carboxyl beads were first modified with an amino polyethylene glycol (PEG) spacer functionalized with an azido group (H_2_N-PEG_23_-N_3_) to provide better display of the sorting signal away from the surface of the bead.^*23*^ The azido group on the bead was then reacted with K(DBCO)LPMTG to install the srtA substrate *via* strain-promoted click chemistry.^*48*^ To ensure that H_2_N-PEG_23_-N_3_ was successfully coupled to the beads, we initially treated some of the beads with DBCO-Fl, washed away excess DBCO-Fl, and monitored the fluorescence of the beads via the use of flow cytometry. An observable increase in fluorescence over beads without H_2_N-PEG_23_-N_3_ was recorded (**Figure S5**), confirming that the beads were displaying N_3_ on the surface.

**Figure 4.**
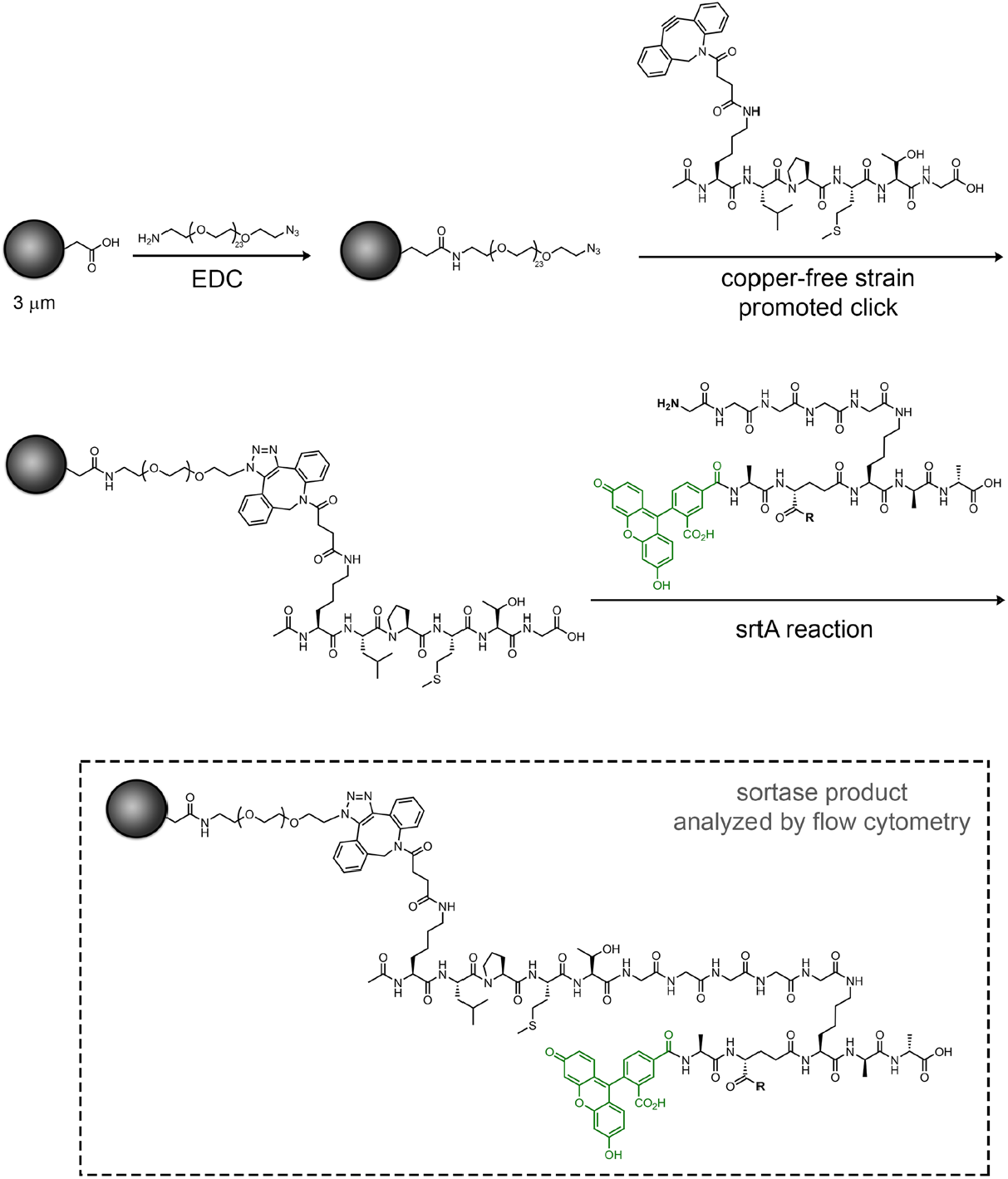
Schematic representation of the sortase-mediated ligation of the fluorescently-linked stem peptide analog onto a bead. The beads are analyzed by flow cytometry and the fluorescence level is reflective of the sortase-related ligation.

With the KLPMTG linked beads in hand, we turned our attention to quantitatively analyze srtA reactions. For all bead-based assays, the stem peptides were modified with fluorescein (fl) instead of TAMRA due to better compatibility for the flow cytometer. Initially, the amidation of D-iGlu was analyzed in the context of a full length pentapeptide in the molecules **flP**^**Q**^**gly5** and **flP**^**E**^**gly5** (**Figure 6A**). Amidation at the carboxyl of the iso-D-glutamic acid residue (which results in iso-D-glutamine) is a PG modification catalyzed by the MurT and GatD enzyme complex.^*49, 50*^ There has been some speculation about the role of amidation for PG assembly. To this end, prior *in vitro* studies using enzymes from *Streptococcus pneumoniae, S. aureus*, Mycobacteria, and *Enterococcus faecalis* suggested that PG crosslinking by PBPs is disrupted in the absence of amidation.^*37, 51-53*^ In fact, the MurT/GatD complex is essential in a number of human pathogens, making it a potential drug target.^*52, 54, 55*^ As expected, due to srtA-mediated ligation of a fluorescently tagged stem peptide to the sorting signal-containing beads, only when sortase was present was a fluorescent readout observed (**Figure 5B**). Interestingly, there was only a small difference in fluorescence levels between **flP**^**Q**^**gly5** and **flP**^**E**^**gly5** in this assay, which suggests that amidation of iso-D-Glu is not playing a large role in controlling acyl-acceptor strand processing when 5 glycines are present on the cross bridge.

**Figure 5.**
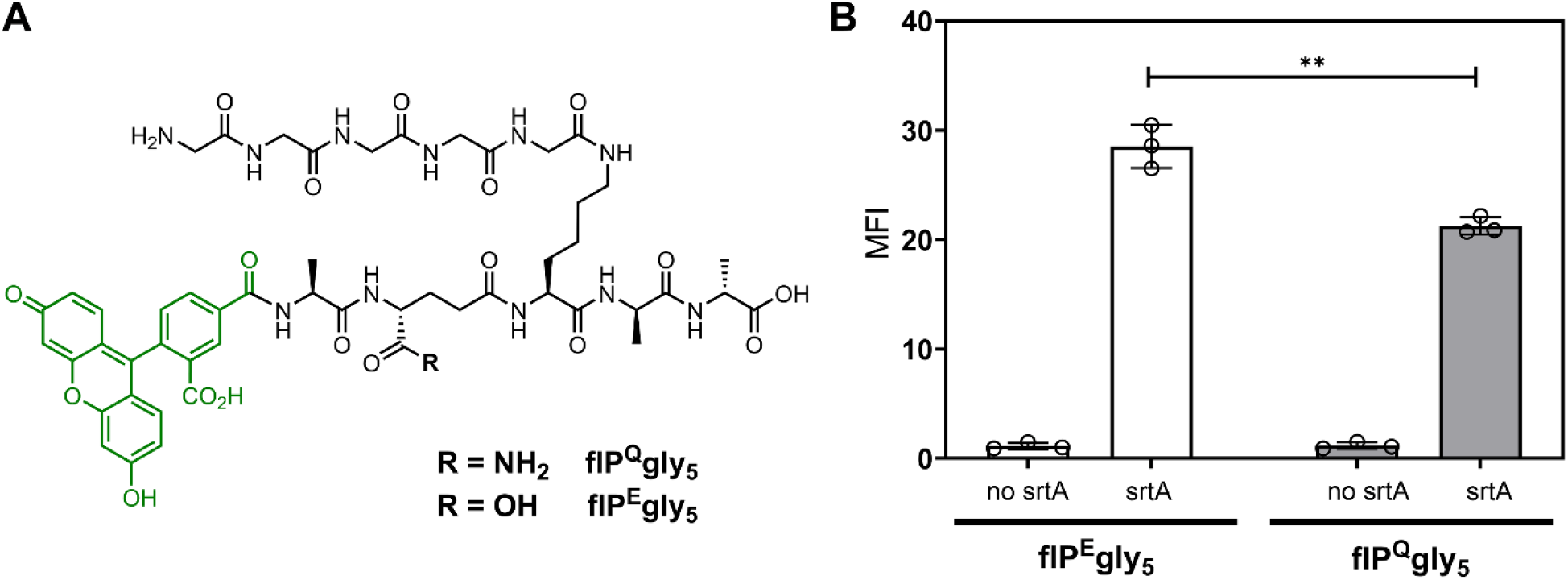
(A) Chemical structures of **flP**^**Q**^**gly**_**5**_ and **flP**^**E**^**gly**_**5**_. (B) Flow cytometry analysis of beads modified with the srtA sorting signal, then treated with stem peptide analogs in the presence or absence of srtA. Data are represented as mean +/-SD (n = 3). *P*-values were determined by a two-tailed *t*-test (* denotes a *p*-value < 0.05, ** < 0.01, ***<0.001, ns = not significant).

**Figure 6.**
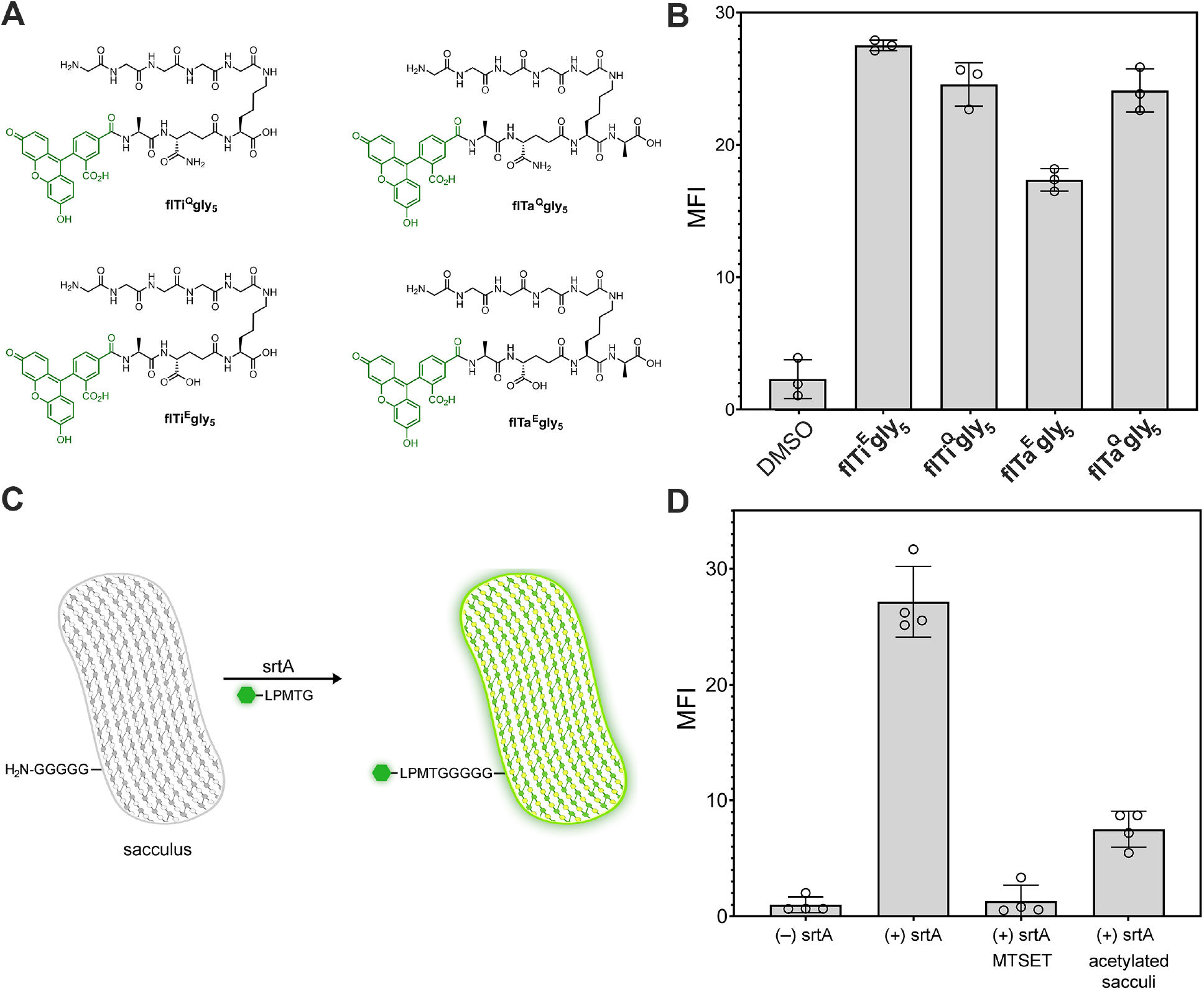
(A) Chemical structure of tripeptide and tetrapeptide analogs of *S. aureus* PG. Each has a fluorescein linked to its *N*-terminus for quantification. (B) Flow cytometry analysis of beads modified with the srtA sorting signal, then treated with various stem peptide analogs in the presence or absence (DMSO) of srtA. Data are represented as mean +/-SD (n = 3). (C) Schematic representation of the SaccuFlow analysis of the srtA processing of the sorting signal modified with a fluorescent handle. (D) Flow cytometry analysis of sacculi incubated with the stated conditions. Data are represented as mean +/-SD (n = 3).

Owing to the fact that it is possible that srtA is operating on stem peptides of various lengths, we turned our attention to shorter versions of the stem peptide (tetrapeptides, tripeptides) that are known to be highly abundant within the PG scaffold of *S. aureus*.^*56-59*^ While tetra- and tripeptides can act as substrates in transpeptidase-catalyzed reactions for PG crosslinking, we wanted to examine the extent to which they participate in sortase A-mediated ligation reactions. As such, we synthesized a series of trimeric and tetrameric stem peptide analogs to examine the extent to which sortase utilizes them as substrates using the bead assay (**Figure 6A**). Our data showed that there is not a large effect in terms of both amidation and stem peptide length. From these results, it appears that srtA does not have a strong preference for a specific length of the stem peptide or amidation of D-iGlu. These results provide evidence that it may be possible for srtA (and other related sortases) to covalently link proteins onto the mature PG, as well as the nascent PG unit. To test this concept, we isolated sacculi from *S. aureus* and incubated it with srtA and a fluorescein-tagged LMPTG sorting signal. Similar to the bead assay, we expect that processing by srtA will result in the covalent linking of LPMTG onto the sacculi, which is analyzed by flow cytometry (**Figure 6C**). The sacculi are devoid of lipid II, and, instead, only will have mature PG. Our results showed that efficient transpeptidation is observed in the absence of lipid II (**Figure 6D**). Inhibition of srtA with methanethiosulfonate (MTSET), a covalent inhibitor of sortase^*60*^ led to a significant decrease in fluorescence levels. Moreover, when the acyl-donor sites are chemically blocked by acetylation, there is a significant drop in fluorescence levels confirming the necessity of free amino groups in the cross-bridge for acylation. Together, these results are consistent with the prospect that processing by srtA may be possible in various PG fragments and may not be restricted to lipid II. In support of this, it was recently shown that incubation of bacterial cells with LPXTG modified probes led to incorporation within the mature PG, not lipid II.^*61*^

## Conclusion

We have assembled a systematic analysis of the possible stem peptides from *S. aureus* that can act as acyl-acceptor strands. These analogs probed the length of the cross-bridge, the amidation state of D-iGlu, and the length of the stem peptide backbone. To quantitatively measure srtA activity, we developed a flow-based bead assay that is robust and has the potential to be compatible with high-throughput analysis. Our results showed that srtA activity was mostly independent of the individual structural modifications we selected. These results could indicate that srtA can, potentially, anchor proteins onto mature PG and does not necessarily exclusively utilize lipid II as the acyl-acceptor strand, as it had been described previously. To the best of our knowledge, this is the first analysis of stem peptide variation in the activity of srtA. We anticipate that a similar platform can be applied to other sortases or enzyme classes (e.g., Braun protein from *Escherichia coli*) to better understand how the primary structure of the PG could potentially modulate its processing by cell wall-linked enzymes.

## References

1. Vollmer, W., Blanot, D., and De Pedro, M. A. (2008) Peptidoglycan structure and architecture, FEMS Microbiology Reviews 32, 149–167. DOI: 10.1111/j.1574-6976.2007.00094.x.

2. Schleifer, K. H., and Kandler, O. (1972) Peptidoglycan types of bacterial cell walls and their taxonomic implications, Bacteriol Rev 36, 407–477. DOI: 10.1128/br.36.4.407-477.1972.

3. Strandén, A. M., Ehlert, K., Labischinski, H., and Berger-Bächi, B. (1997) Cell wall monoglycine cross-bridges and methicillin hypersusceptibility in a femAB null mutant of methicillin-resistant Staphylococcus aureus, Journal of Bacteriology 179, 9–16. DOI: 10.1128/jb.179.1.9-16.1997.

4. Foster, T. J., Geoghegan, J. A., Ganesh, V. K., and Hook, M. (2014) Adhesion, invasion and evasion: the many functions of the surface proteins of Staphylococcus aureus, Nat Rev Microbiol 12, 49–62. DOI: 10.1038/nrmicro3161.

5. Uhlén, M., Guss, B., Nilsson, B., Götz, F., and Lindberg, M. (1984) Expression of the gene encoding protein A in Staphylococcus aureus and coagulase-negative staphylococci, J Bacteriol 159, 713–719. DOI: 10.1128/jb.159.2.713-719.1984.

6. Sjodahl, J. (1977) Repetitive sequences in protein A from Staphylococcus aureus. Arrangement of five regions within the protein, four being highly homologous and Fc-binding, Eur J Biochem 73, 343–351. DOI: 10.1111/j.1432-1033.1977.tb11324.x.

7. Jensen, K. (2007) A normally occurring Staphylococcus antibody in human serum, Apmis 115, 533–539; discussion 540-531. DOI: 10.1111/j.1600-0463.2007.apm_731a.x.

8. Mazmanian, S. K., Liu, G., Ton-That, H., and Schneewind, O. (1999) <em>Staphylococcus aureus</em> Sortase, an Enzyme that Anchors Surface Proteins to the Cell Wall, Science 285, 760–763. DOI: 10.1126/science.285.5428.760.

9. Ton-That, H., Liu, G., Mazmanian, S. K., Faull, K. F., and Schneewind, O. (1999) Purification and characterization of sortase, the transpeptidase that cleaves surface proteins of <em>Staphylococcus aureus</em> at the LPXTG motif, Proceedings of the National Academy of Sciences 96, 12424–12429. DOI: 10.1073/pnas.96.22.12424.

10. Mazmanian, S. K., Ton-That, H., and Schneewind, O. (2001) Sortase-catalysed anchoring of surface proteins to the cell wall of Staphylococcus aureus, Mol Microbiol 40, 1049–1057. DOI: 10.1046/j.1365-2958.2001.02411.x.

11. Marraffini, L. A., Ton-That, H., Zong, Y., Narayana, S. V. L., and Schneewind, O. (2004) Anchoring of Surface Proteins to the Cell Wall of <em>Staphylococcus aureus</em>: A CONSERVED ARGININE RESIDUE IS REQUIRED FOR EFFICIENT CATALYSIS OF SORTASE A *, Journal of Biological Chemistry 279, 37763–37770. DOI: 10.1074/jbc.M405282200.

12. Aulabaugh, A., Ding, W., Kapoor, B., Tabei, K., Alksne, L., Dushin, R., Zatz, T., Ellestad, G., and Huang, X. (2007) Development of an HPLC assay for Staphylococcus aureus sortase: evidence for the formation of the kinetically competent acyl enzyme intermediate, Anal Biochem 360, 14–22. DOI: 10.1016/j.ab.2006.10.021.

13. Ton-That, H., Labischinski, H., Berger-Bächi, B., and Schneewind, O. (1998) Anchor Structure of Staphylococcal Surface Proteins: III. ROLE OF THE FemA, FemB, AND FemX FACTORS IN ANCHORING SURFACE PROTEINS TO THE BACTERIAL CELL WALL*, Journal of Biological Chemistry 273, 29143–29149. DOI: https://doi.org/10.1074/jbc.273.44.29143.

14. Perry, A. M., Ton-That, H., Mazmanian, S. K., and Schneewind, O. (2002) Anchoring of surface proteins to the cell wall of Staphylococcus aureus. III. Lipid II is an in vivo peptidoglycan substrate for sortase-catalyzed surface protein anchoring, J Biol Chem 277, 16241–16248. DOI: 10.1074/jbc.M109194200.

15. Ruzin, A., Severin, A., Ritacco, F., Tabei, K., Singh, G., Bradford, P. A., Siegel, M. M., Projan, S. J., and Shlaes, D. M. (2002) Further Evidence that a Cell Wall Precursor [C<sub>55</sub>-MurNAc-(Peptide)-GlcNAc] Serves as an Acceptor in a Sorting Reaction, Journal of Bacteriology 184, 2141–2147. DOI: 10.1128/jb.184.8.2141-2147.2002.

16. Mazmanian, S. K., Liu, G., Jensen, E. R., Lenoy, E., and Schneewind, O. (2000) Staphylococcus aureus sortase mutants defective in the display of surface proteins and in the pathogenesis of animal infections, Proc Natl Acad Sci U S A 97, 5510–5515. DOI: 10.1073/pnas.080520697.

17. Apostolos, A. J., Ferraro, N. J., Dalesandro, B. E., and Pires, M. M. (2021) SaccuFlow: A High-Throughput Analysis Platform to Investigate Bacterial Cell Wall Interactions, ACS Infect Dis 7, 2483–2491. DOI: 10.1021/acsinfecdis.1c00255.

18. Cascioferro, S., Totsika, M., and Schillaci, D. (2014) Sortase A: an ideal target for anti-virulence drug development, Microb Pathog 77, 105–112. DOI: 10.1016/j.micpath.2014.10.007.

19. Zhang, J., Liu, H., Zhu, K., Gong, S., Dramsi, S., Wang, Y. T., Li, J., Chen, F., Zhang, R., Zhou, L., Lan, L., Jiang, H., Schneewind, O., Luo, C., and Yang, C. G. (2014) Antiinfective therapy with a small molecule inhibitor of Staphylococcus aureus sortase, Proc Natl Acad Sci U S A 111, 13517–13522. DOI: 10.1073/pnas.1408601111.

20. Popp, M. W., and Ploegh, H. L. (2011) Making and breaking peptide bonds: protein engineering using sortase, Angew Chem Int Ed Engl 50, 5024–5032. DOI: 10.1002/anie.201008267.

21. Kruger, R. G., Dostal, P., and McCafferty, D. G. (2002) An economical and preparative orthogonal solid phase synthesis of fluorescein and rhodamine derivatized peptides: FRET substrates for the Staphylococcus aureus sortase SrtA transpeptidase reaction, Chem Commun (Camb), 2092–2093. DOI: 10.1039/b206303d.

22. Mao, H., Hart, S. A., Schink, A., and Pollok, B. A. (2004) Sortase-mediated protein ligation: a new method for protein engineering, J Am Chem Soc 126, 2670–2671. DOI: 10.1021/ja039915e.

23. Parthasarathy, R., Subramanian, S., and Boder, E. T. (2007) Sortase A as a Novel Molecular “Stapler” for Sequence-Specific Protein Conjugation, Bioconjugate Chemistry 18, 469–476. DOI: 10.1021/bc060339w.

24. Glasgow, J. E., Salit, M. L., and Cochran, J. R. (2016) In Vivo Site-Specific Protein Tagging with Diverse Amines Using an Engineered Sortase Variant, Journal of the American Chemical Society 138, 7496–7499. DOI: 10.1021/jacs.6b03836.

25. Lavollay, M., Arthur, M., Fourgeaud, M., Dubost, L., Marie, A., Veziris, N., Blanot, D., Gutmann, L., and Mainardi, J.-L. (2008) The Peptidoglycan of Stationary-Phase <em>Mycobacterium tuberculosis</em> Predominantly Contains Cross-Links Generated by <span class=“sc”>l,d</span>-Transpeptidation, Journal of Bacteriology 190, 4360–4366. DOI: 10.1128/jb.00239-08.

26. Spratt, B. G., and Pardee, A. B. (1975) Penicillin-binding proteins and cell shape in E. coli, Nature 254, 516–517. DOI: 10.1038/254516a0.

27. Monteiro, J. M., Covas, G., Rausch, D., Filipe, S. R., Schneider, T., Sahl, H. G., and Pinho, M. G. (2019) The pentaglycine bridges of Staphylococcus aureus peptidoglycan are essential for cell integrity, Sci Rep 9, 5010. DOI: 10.1038/s41598-019-41461-1.

28. Roemer, T., Schneider, T., and Pinho, M. G. (2013) Auxiliary factors: a chink in the armor of MRSA resistance to beta-lactam antibiotics, Curr Opin Microbiol 16, 538–548. DOI: 10.1016/j.mib.2013.06.012.

29. Figueiredo, T. A., Sobral, R. G., Ludovice, A. M., Almeida, J. M., Bui, N. K., Vollmer, W., de Lencastre, H., and Tomasz, A. (2012) Identification of genetic determinants and enzymes involved with the amidation of glutamic acid residues in the peptidoglycan of Staphylococcus aureus, PLoS Pathog 8, e1002508. DOI: 10.1371/journal.ppat.1002508.

30. Munch, D., Roemer, T., Lee, S. H., Engeser, M., Sahl, H. G., and Schneider, T. (2012) Identification and in vitro analysis of the GatD/MurT enzyme-complex catalyzing lipid II amidation in Staphylococcus aureus, PLoS Pathog 8, e1002509. DOI: 10.1371/journal.ppat.1002509.

31. Antos, J. M., Chew, G. L., Guimaraes, C. P., Yoder, N. C., Grotenbreg, G. M., Popp, M. W., and Ploegh, H. L. (2009) Site-specific N-and C-terminal labeling of a single polypeptide using sortases of different specificity, J Am Chem Soc 131, 10800–10801. DOI: 10.1021/ja902681k.

32. Antos, J. M., Miller, G. M., Grotenbreg, G. M., and Ploegh, H. L. (2008) Lipid modification of proteins through sortase-catalyzed transpeptidation, J Am Chem Soc 130, 16338–16343. DOI: 10.1021/ja806779e.

33. Guimaraes, C. P., Witte, M. D., Theile, C. S., Bozkurt, G., Kundrat, L., Blom, A. E., and Ploegh, H. L. (2013) Site-specific C-terminal and internal loop labeling of proteins using sortase-mediated reactions, Nat Protoc 8, 1787–1799. DOI: 10.1038/nprot.2013.101.

34. Chen, L., Cohen, J., Song, X., Zhao, A., Ye, Z., Feulner, C. J., Doonan, P., Somers, W., Lin, L., and Chen, P. R. (2016) Improved variants of SrtA for site-specific conjugation on antibodies and proteins with high efficiency, Sci Rep 6, 31899–31899. DOI: 10.1038/srep31899.

35. Guimaraes, C. P., Witte, M. D., Theile, C. S., Bozkurt, G., Kundrat, L., Blom, A. E. M., and Ploegh, H. L. (2013) Site-specific C-terminal and internal loop labeling of proteins using sortase-mediated reactions, Nature Protocols 8, 1787–1799. DOI: 10.1038/nprot.2013.101.

36. Popp, M. W., Dougan, S. K., Chuang, T.-Y., Spooner, E., and Ploegh, H. L. (2011) Sortase-catalyzed transformations that improve the properties of cytokines, Proceedings of the National Academy of Sciences 108, 3169–3174. DOI: 10.1073/pnas.1016863108.

37. Pidgeon, S. E., Apostolos, A. J., Nelson, J. M., Shaku, M., Rimal, B., Islam, M. N., Crick, D. C., Kim, S. J., Pavelka, M. S., Kana, B. D., and Pires, M. M. (2019) L,D-Transpeptidase Specific Probe Reveals Spatial Activity of Peptidoglycan Cross-Linking, ACS Chem Biol 14, 2185–2196. DOI: 10.1021/acschembio.9b00427.

38. Dalesandro, B. E., and Pires, M. M. (2021) Induction of Endogenous Antibody Recruitment to the Surface of the Pathogen Enterococcus faecium, ACS Infect Dis 7, 1116–1125. DOI: 10.1021/acsinfecdis.0c00547.

39. Apostolos, A. J., Pidgeon, S. E., and Pires, M. M. (2020) Remodeling of Cross-bridges Controls Peptidoglycan Cross-linking Levels in Bacterial Cell Walls, ACS Chem Biol 15, 1261–1267. DOI: 10.1021/acschembio.0c00002.

40. Apostolos, A. J., Nelson, J. M., Silva, J. R. A., Lameira, J., Achimovich, A. M., Gahlmann, A., Alves, C. N., and Pires, M. M. (2020) Facile Synthesis and Metabolic Incorporation of m-DAP Bioisosteres Into Cell Walls of Live Bacteria, ACS Chem Biol 15, 2966–2975. DOI: 10.1021/acschembio.0c00618.

41. Ngadjeua, F., Braud, E., Saidjalolov, S., Iannazzo, L., Schnappinger, D., Ehrt, S., Hugonnet, J. E., Mengin-Lecreulx, D., Patin, D., Etheve-Quelquejeu, M., Fonvielle, M., and Arthur, M. (2018) Critical Impact of Peptidoglycan Precursor Amidation on the Activity of l,d-Transpeptidases from Enterococcus faecium and Mycobacterium tuberculosis, Chemistry 24, 5743–5747. DOI: 10.1002/chem.201706082.

42. Gautam, S., Kim, T., Shoda, T., Sen, S., Deep, D., Luthra, R., Ferreira, M. T., Pinho, M. G., and Spiegel, D. A. (2015) An Activity-Based Probe for Studying Crosslinking in Live Bacteria, Angew Chem Int Ed Engl 54, 10492–10496. DOI: 10.1002/anie.201503869.

43. Lin, H., Lin, L., Du, Y., Gao, J., Yang, C., and Wang, W. (2021) Biodistributions of l,d-Transpeptidases in Gut Microbiota Revealed by In Vivo Labeling with Peptidoglycan Analogs, ACS Chem Biol 16, 1164–1171. DOI: 10.1021/acschembio.1c00346.

44. Ehlert, K., Schröder, W., and Labischinski, H. (1997) Specificities of FemA and FemB for different glycine residues: FemB cannot substitute for FemA in staphylococcal peptidoglycan pentaglycine side chain formation, Journal of Bacteriology 179, 7573–7576. DOI: 10.1128/jb.179.23.7573-7576.1997.

45. de Jonge, B. L., Sidow, T., Chang, Y. S., Labischinski, H., Berger-Bachi, B., Gage, D. A., and Tomasz, A. (1993) Altered muropeptide composition in Staphylococcus aureus strains with an inactivated femA locus, Journal of Bacteriology 175, 2779–2782. DOI: 10.1128/jb.175.9.2779-2782.1993.

46. Henze, U., Sidow, T., Wecke, J., Labischinski, H., and Berger-Bächi, B. (1993) Influence of femB on methicillin resistance and peptidoglycan metabolism in Staphylococcus aureus, Journal of Bacteriology 175, 1612–1620. DOI: 10.1128/jb.175.6.1612-1620.1993.

47. Ton-That, H., Labischinski, H., Berger-Bachi, B., and Schneewind, O. (1998) Anchor structure of staphylococcal surface proteins. III. Role of the FemA, FemB, and FemX factors in anchoring surface proteins to the bacterial cell wall, J Biol Chem 273, 29143–29149. DOI: 10.1074/jbc.273.44.29143.

48. Jewett, J. C., Sletten, E. M., and Bertozzi, C. R. (2010) Rapid Cu-Free Click Chemistry with Readily Synthesized Biarylazacyclooctynones, Journal of the American Chemical Society 132, 3688–3690. DOI: 10.1021/ja100014q.

49. Münch, D., Roemer, T., Lee, S. H., Engeser, M., Sahl, H. G., and Schneider, T. (2012) Identification and in vitro Analysis of the GatD/MurT Enzyme-Complex Catalyzing Lipid II Amidation in Staphylococcus aureus, PLOS Pathogens 8, e1002509. DOI: 10.1371/journal.ppat.1002509.

50. Figueiredo, T. A., Sobral, R. G., Ludovice, A. M., de Almeida, J. M. F., Bui, N. K., Vollmer, W., de Lencastre, H., and Tomasz, A. (2012) Identification of Genetic Determinants and Enzymes Involved with the Amidation of Glutamic Acid Residues in the Peptidoglycan of Staphylococcus aureus, PLOS Pathogens 8, e1002508. DOI: 10.1371/journal.ppat.1002508.

51. Welsh, M. A., Taguchi, A., Schaefer, K., Van Tyne, D., Lebreton, F., Gilmore, M. S., Kahne, D., and Walker, S. (2017) Identification of a Functionally Unique Family of Penicillin-Binding Proteins, J Am Chem Soc 139, 17727–17730. DOI: 10.1021/jacs.7b10170.

52. Zapun, A., Philippe, J., Abrahams, K. A., Signor, L., Roper, D. I., Breukink, E., and Vernet, T. (2013) In vitro reconstitution of peptidoglycan assembly from the * Gram-positive pathogen Streptococcus pneumoniae, ACS Chem Biol 8, 2688–2696. DOI: 10.1021/cb400575t.

53. Squeglia, F., Ruggiero, A., and Berisio, R. (2018) Chemistry of Peptidoglycan in Mycobacterium tuberculosis Life Cycle: An off-the-wall Balance of Synthesis and Degradation, Chemistry 24, 2533–2546. DOI: 10.1002/chem.201702973.

54. DeJesus, M. A., Gerrick, E. R., Xu, W., Park, S. W., Long, J. E., Boutte, C. C., Rubin, E. J., Schnappinger, D., Ehrt, S., Fortune, S. M., Sassetti, C. M., and Ioerger, T. R. (2017) Comprehensive Essentiality Analysis of the Mycobacterium tuberculosis Genome via Saturating Transposon Mutagenesis, mBio 8. DOI: 10.1128/mBio.02133-16.

55. Liu, X., Gallay, C., Kjos, M., Domenech, A., Slager, J., van Kessel, S. P., Knoops, K., Sorg, R. A., Zhang, J. R., and Veening, J. W. (2017) High-throughput CRISPRi phenotyping identifies new essential genes in Streptococcus pneumoniae, Mol Syst Biol 13, 931. DOI: 10.15252/msb.20167449.

56. Typas, A., Banzhaf, M., Gross, C. A., and Vollmer, W. (2011) From the regulation of peptidoglycan synthesis to bacterial growth and morphology, Nat Rev Microbiol 10, 123–136. DOI: 10.1038/nrmicro2677.

57. Snowden, M. A., and Perkins, H. R. (1990) Peptidoglycan cross-linking in Staphylococcus aureus. An apparent random polymerisation process, Eur J Biochem 191, 373–377. DOI: 10.1111/j.1432-1033.1990.tb19132.x.

58. Snowden, M. A., and Perkins, H. R. (1991) Cross-linking and O-acetylation of peptidoglycan in Staphylococcus aureus (strains H and MR-1) grown in the * presence of sub-growth-inhibitory concentrations of beta-lactam antibiotics, J Gen Microbiol 137, 1661–1666. DOI: 10.1099/00221287-137-7-1661.

59. Sutton, J. A. F., Carnell, O. T., Lafage, L., Gray, J., Biboy, J., Gibson, J. F., Pollitt, E. J. G., Tazoll, S. C., Turnbull, W., Hajdamowicz, N. H., Salamaga, B., Pidwill, G. R., Condliffe, A. M., Renshaw, S. A., Vollmer, W., and Foster, S. J. (2021) Staphylococcus aureus cell wall structure and dynamics during host-pathogen interaction, PLoS Pathog 17, e1009468. DOI: 10.1371/journal.ppat.1009468.

60. Ton-That, H., and Schneewind, O. (1999) Anchor structure of staphylococcal surface proteins. IV. Inhibitors of the cell wall sorting reaction, J Biol Chem 274, 24316–24320. DOI: 10.1074/jbc.274.34.24316.

61. Hansenova Manaskova, S., Nazmi, K., Van’t Hof, W., van Belkum, A., Kaman, W. E., Martin, N. I., Veerman, E. C. I., and Bikker, F. J. (2022) Natural and Synthetic Sortase A Substrates Are Processed by Staphylococcus aureus via Different Pathways, Bioconjug Chem 33, 555–559. DOI: 10.1021/acs.bioconjchem.2c00012.

