## Supporting Information for "Structure Activity Relationship of the Stem Peptide in Sortase A mediated Ligation from *Staphylococcus aureus*"

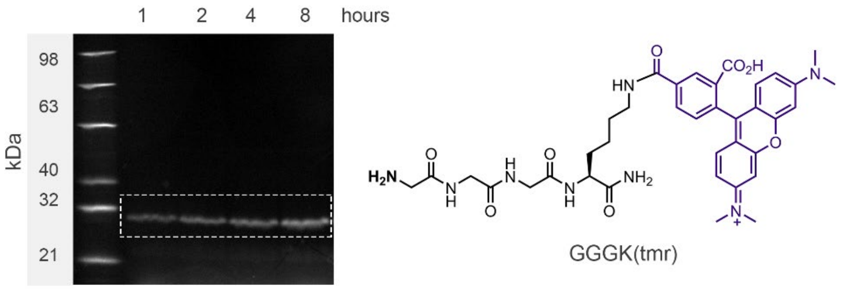


**Figure S1.** Gel (SDS-PAGE) fluorescence analysis of time points of GGGK(tmr) where the reaction with all elements was quenched at 1, 2, 4, 8 h and displayed an increase in fluorescence as time progressed. An expected product band of ~29 kDa was anticipated for GFP-LPETGGGK(tmr) products.

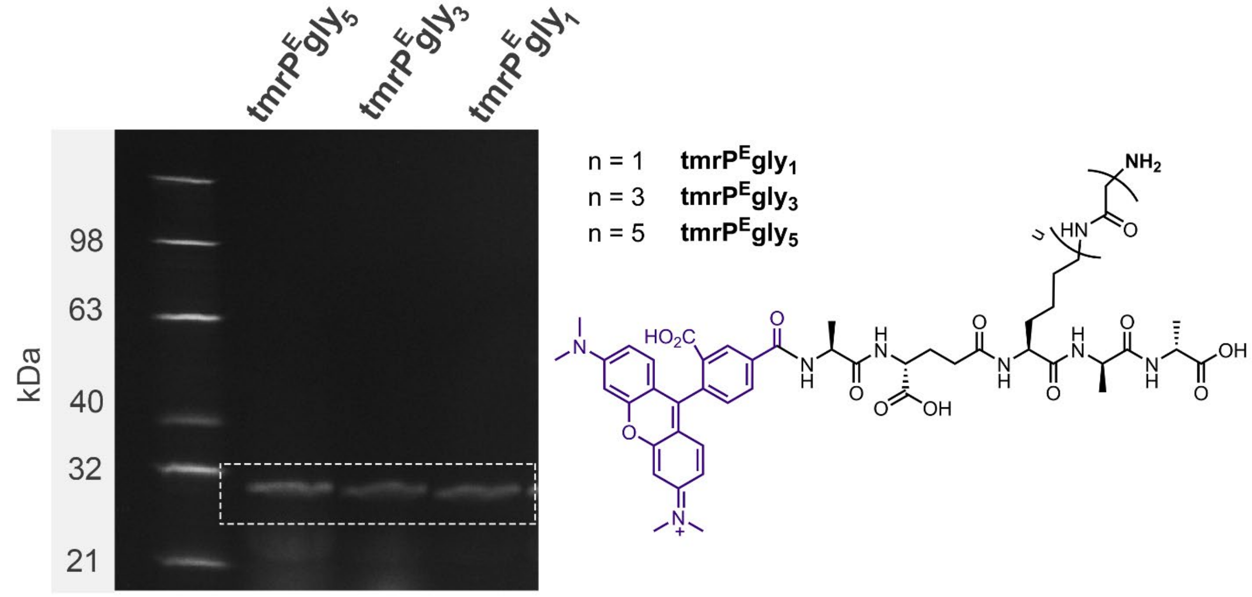


**Figure S2.** Gel (SDS-PAGE) fluorescence analysis of **tmrP$^{E}$gly$_5$**, **tmrP$^{E}$gly$_3$**, and **tmrP$^{E}$gly$_1$** after incubation of reaction elements for 8 h. Product band is calculated to be ~29 kDa.

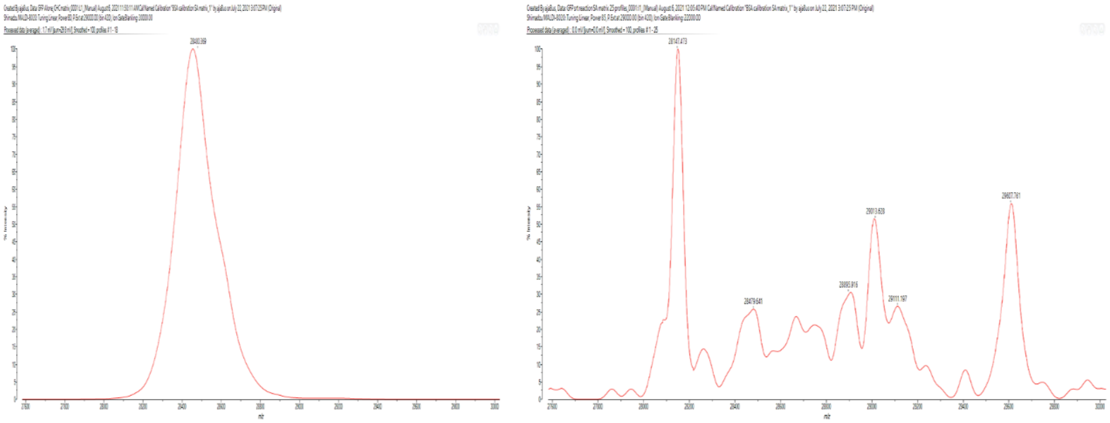


**Figure S3.** MALDI-TOF analysis of GFP-LPETG alone (left) and GFP-LPETG incubated with all reaction elements and **tmrP^E^gly$_5$** (right).

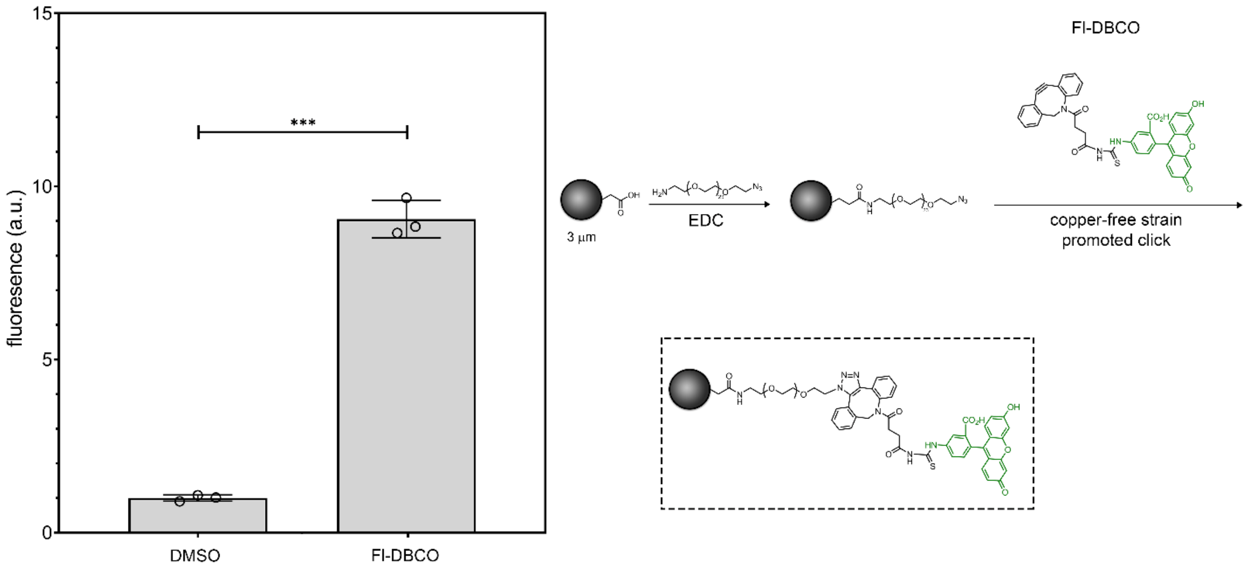


**Figure S4.** Flow cytometry analysis of $H_2N$-$PEG_{23}$-$N_3$ functionalized beads treated with DBCO-Fl (25 mm) for 30 min at RT. Beads were washed 3X with 1X PBS before analysis. Data are represented as mean +/- SD (n = 3). *P*-values were determined by a two-tailed *t*-test (* denotes a *p*-value < 0.05, ** < 0.01, ***<0.001, ns = not significant).

**Materials.** All peptide related reagents (resin, coupling reagent, deprotection reagent, amino acids, and cleavage reagents) were purchased from ChemImpex. Carboxyl polystyrene beads were purchased from Spherotech. Azido-Peg23-Amine was purchased from Broadpharm. Precast gels (10-20% Tricine 1.0 mm) were purchased from Invitrogen.

**SrtA in-gel fluorescence Assays.** Enzymatic reactions were set up with $V_F$ = 110 μL. In each reaction, 10 μM GFP-LPETG, 10 μM SrtA, 500 μM respective (tmr)peptide, 1X sortase buffer (10X contains 500 mM Tris-HCl, pH 7.5, 1.5 M NaCl, 100 mM CaCl2) were incubated for the time points described. Reactions were quenched with 0.1% TFA. The plasmid for sortase A from *S. aureus* was obtained from Addgene: pET28a-SrtAdelta59 (plasmid no. 51138). The plasmid for GFP-LPETG was also obtained from Addgene (plasmid no. 71754). Each protein was expressed and purified as described by the depositing labs. Invitrogen Novex 10 to 20% Tricine protein precast gels were used to analyze srtA reactions. Each well was loaded with 15 μL of sample and run at 120 V for 75 min. The fluorescent protein standard was purchased from Invitrogen (Cat. No. 10747012) and 5 μL was loaded in each gel. The gel was imaged using a BioRad ChemiDoc XRS+ imager equipped with a XcitaBlue conversion screen with exposure time set to 15 s for all images obtained.

**SrtA SPHERO Carboxy Polystyrene Particles Assays.** Sphero Carboxy Polystryene beads 3.31 um 5% w/v (Spherotech, 250 μL) were reacted with 0.05 M MES at pH 6 (250 μL), EDC (10 mg), and $NH_2$-Peg23-Az (BroadPharm, 2 mg) for 24 h. Beads were collected at 3,000g for 15 min. The supernatant was carefully removed, and the beads were washed in PBS. A final working suspension of 5% w/v was made for assays. K(DBCO)LPMTG was reacted with the beads displaying Peg23-Az once the beads were blocked with Tween 80 (0.05%) in PBS, washed, and then resuspended in PBS. The particles were washed as before and resuspended to a 5% w/v suspension. SrtA (10 μM) was incubated with the beads (2.5 μL), 100 μM of each (fl)peptide, and 1X sortase buffer for 4 h at room temperature before the reaction was quenched with 0.1% TFA and washed 3X with 10% SDS. Samples were then diluted 2-fold in PBS and analyzed using an Attune NxT flow cytometer equipped with a 488 nm laser and 525/40 nm bandpass filter. The data were analyzed using the Attune NxT Software, where populations were gated and no less than 10,000 events per sample were recorded.

**SrtA Enzymatic Assay with Sacculi**. Isolated *S. aureus* sacculi samples were incubated with 20 uM sortase A, 100 uM sorting signal substrate (FL-LPMTG), and 1X sortase buffer (10X contains 500 mM Tris-HCl, pH 7.5, 1.5 M NaCl, 100 mM CaCl2). Samples that contained the covalent inhibitor, MTSET, were run at a concentration of 1 mM. Sacculi (1.6 eq.) that was acetylated was treated with acetic anhydride (1 eq.), DIEA (1.7 eq), in DMF (10 eq.) for 1 h at room temp. Sacculi was then harvested and resuspended in PBS. All samples were shook at room temperature for 4 h, quenched with 0.1% TFA, and washed 3X with freshly made 8 M urea. Samples were resuspended in a final volume of 200 uL 1X PBS and analyzed by flow cytometry as described above.

**Scheme S1. Synthesis of Tripeptides**

Fmoc-Lys(Mtt)-Wang resin (250 mg) was added to a 25 mL peptide synthesis vessel with 20% piperidine in DMF for 30 min to remove the base-labile protecting group. The resin was washed with MeOH and DCM (3 x 15 mL each). Fmoc-D-glutamic acid α-amide (3 eq, 193 mg, 2.1 mmol) for "Q" / "Gln" peptides *or* Fmoc-D-glutamic acid-OtBu-OH (3 eq, 223 mg, 2.1 mmol) for "E" / "Glu" peptides, HBTU (3 eq, 200 mg, 2.1 mmol), and DIEA (6 eq, 0.183 mL, 4.2 mmol) in DMF (10 mL) were added to the reaction flask and agitated for 2 h at ambient temperature. The resin was washed as before and the Fmoc deprotection and coupling procedure was repeated as before using the same equivalencies with Fmoc-L-Alanine-OH. The Fmoc group of L-alanine was deprotected and coupled with 5(6)-carboxyfluorescein (2 eq, 132 mg, 1.4 mmol) for "FL" peptides or 5(6)-carboxytetramethylrhodamine (2 eq, 151 mg, 1.4 mmol) for "TAMRA" peptides, HBTU (2 eq, 133 mg, 1.4 mmol) and DIEA (4 eq, 0.122 mL, 2.8 mmol) in DMF (10 mL) shaking overnight. The resin was washed as before and the Mtt group of Lys was removed with 15 mL of a 1% TFA solution in DCM with shaking at ambient temperature for 20 min. This step was repeated until a total of 50 mL of 1% TFA was used. Fmoc-Gly-OH (3 eq, 156 mg, 2.1 mmol) was added to the resin with shaking at room temperature for 2 h. The Fmoc group was removed, and this coupling step was repeated 4X to yield a total of 5 Gly on the Lys side chain. The resin was washed as before and added to a solution of TFA/$H_2O$/TIPS (95%, 2.5%, 2.5%, 20 mL) with agitation for 2 h at ambient temperature. The resin was filtered and resulting solution concentrated *in vacuo*. The residue was trituated with cold diethyl ether.

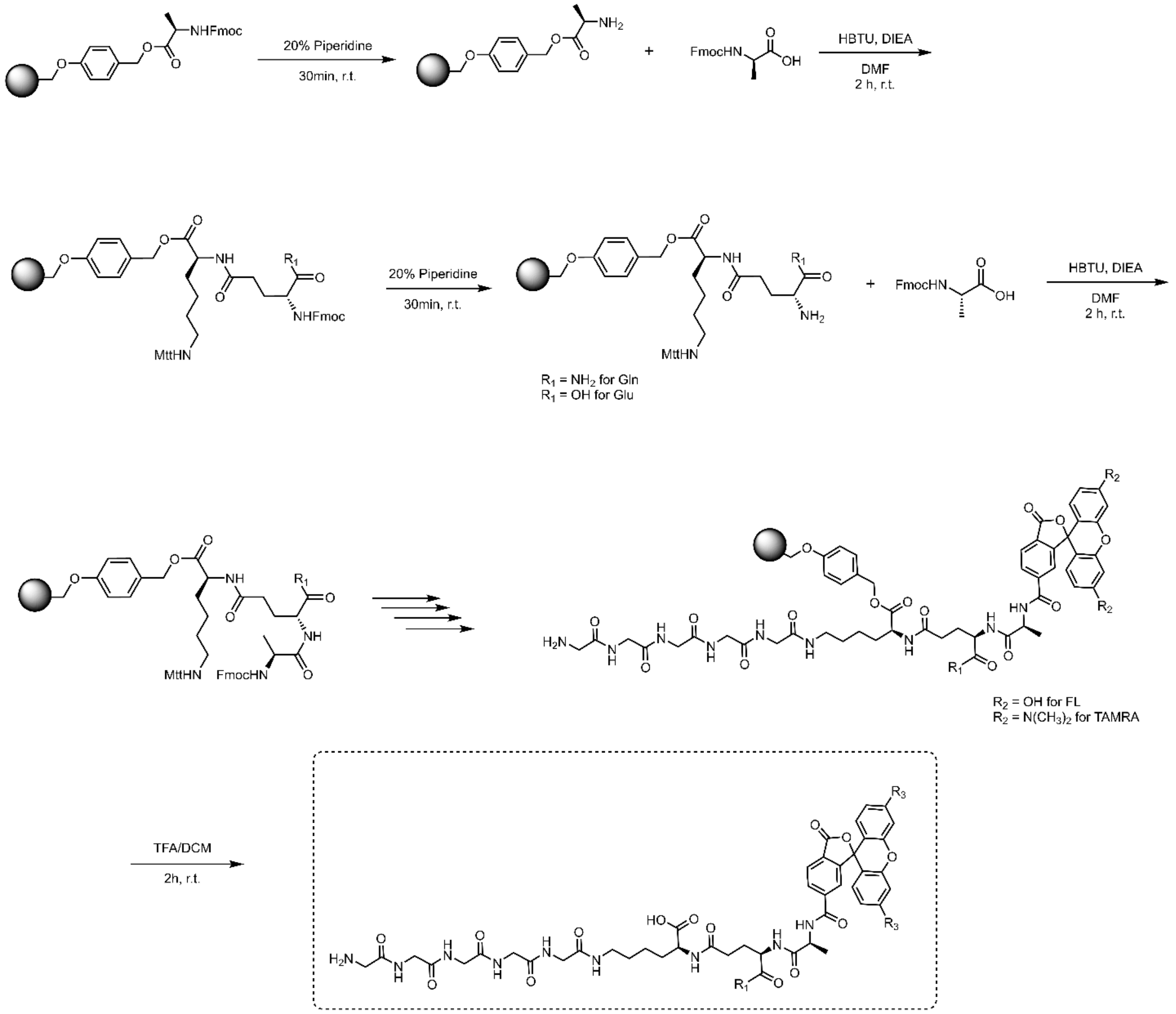

**Scheme S2. Synthesis of Tetrapeptides:**

Fmoc-D-Ala-Wang resin (250 mg) was added to a 25 mL peptide synthesis vessel with 20% piperidine in DMF for 30 min to remove the base-labile protecting group. The resin was washed with MeOH and DCM (3 x 15 mL each). Fmoc-L-Lys(Mtt)-OH (3 eq, 328 mg, 2.1 mmol), HBTU (3 eq, 200 mg, 2.1 mmol), and DIEA (6 eq, 0.183 mL, 4.2 mmol) in DMF (10 mL) were added to the reaction flask and agitated for 2 h at ambient temperature. The resin was washed as before, Fmoc deprotected, and Fmoc-D-glutamic acid α-amide (3 eq, 223 mg, 2.1 mmol) for "Q" / "Gln" peptides *or* Fmoc-D-glutamic acid-OtBu-OH (3 eq, 223 mg, 2.1 mmol) for "E" / "Glu" peptides was added with the same coupling reagents as previously described. The Fmoc deprotection and coupling procedure was repeated as before using the same equivalencies with Fmoc-L-Alanine-OH. The Fmoc group of L-alanine was deprotected and coupled with 5(6)-carboxyfluorescein (2 eq, 132 mg, 1.4 mmol) for "FL" peptides or 5(6)-carboxytetramethylrhodamine (2 eq, 151 mg, 1.4 mmol) for "TAMRA" peptides, HBTU (2 eq, 133 mg, 1.4 mmol) and DIEA (4 eq, 0.122 mL, 2.8 mmol) in DMF (10 mL) shaking overnight. The resin was washed as before and the Mtt group of Lys was removed with 15 mL of a 1% TFA solution in DCM with shaking at ambient temperature for 20 min. This step was repeated until a total of 50 mL of 1% TFA was used. Fmoc-Gly-OH (3 eq, 156 mg, 2.1 mmol) was added to the resin with shaking at room temperature for 2 h. The Fmoc group was removed, and this coupling step was repeated 4X to yield a total of 5 Gly on the Lys side chain. The resin was washed as before and added to a solution of TFA/$H_2O$/TIPS (95%, 2.5%, 2.5%, 20 mL) with agitation for 2 h at ambient temperature. The resin was filtered and resulting solution concentrated *in vacuo*. The residue was trituated with cold diethyl ether.

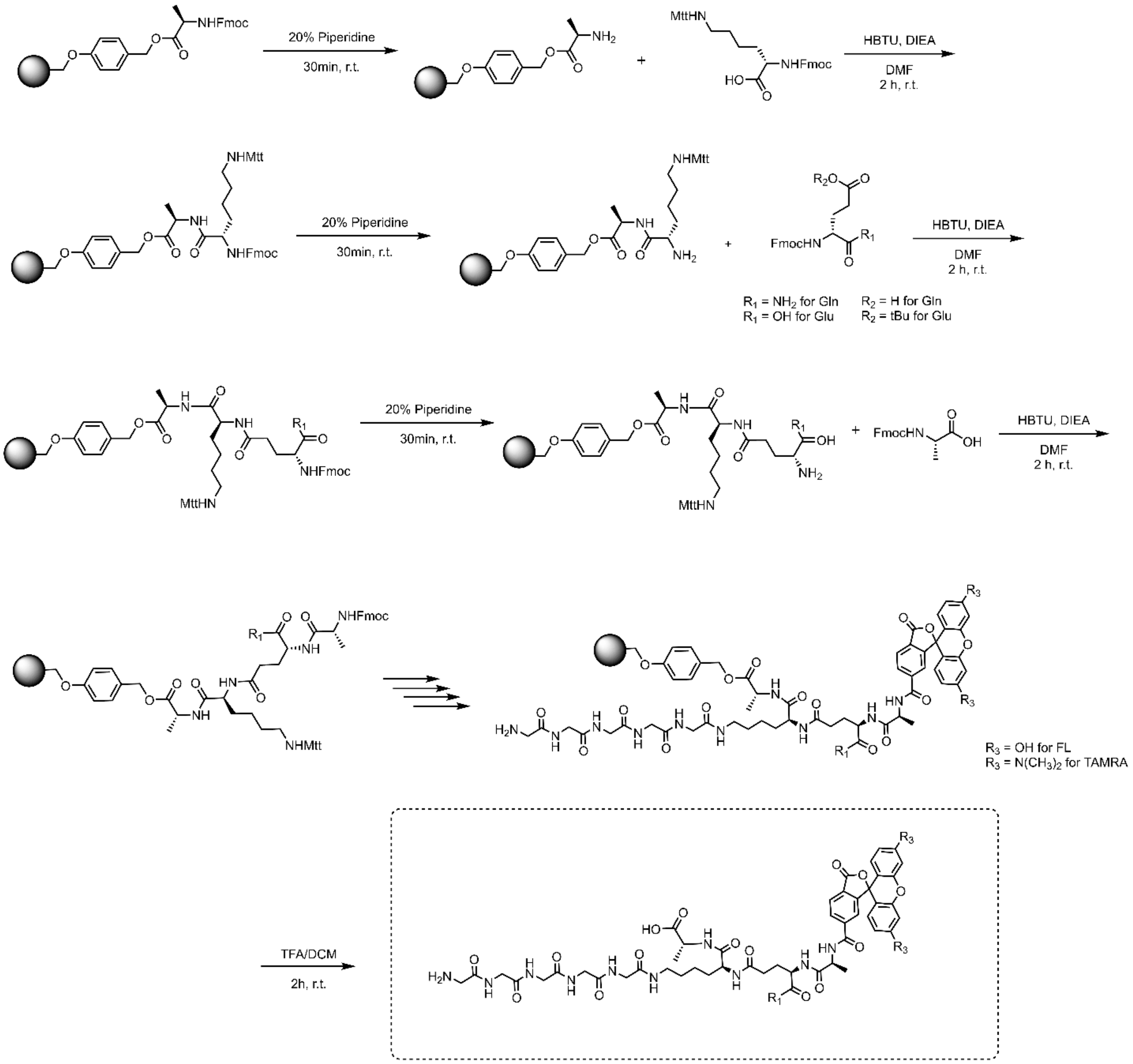
20% Piperidine
30min, r.t.
HBTU, DIEA
DMF
2 h, r.t.
R$_1$ = NH$_2$ for Gln
R$_1$ = OH for Glu
R$_2$ = H for Gln
R$_2$ = tBu for Glu
R$_3$ = OH for FL
R$_3$ = N(CH$_3$)$_2$ for TAMRA
TFA/DCM
2h, r.t.

**Scheme S3. Synthesis of Pentapeptides:**

Fmoc-D-Ala-Wang resin was added to a 25 mL peptide synthesis vessel with 20% piperidine in DMF for 30 min to remove the base-labile protecting group. The resin was washed with MeOH and DCM (3 x 15 mL each). Fmoc-D-Ala-OH (3 eq, 163 mg, 2.1 mmol), HBTU (3 eq, 200 mg, 2.1 mmol), and DIEA (6 eq, 0.183 mL, 4.2 mmol) in DMF (10 mL) were added to the reaction flask and agitated for 2 h at ambient temperature. The resin was washed, Fmoc-deprotected, and Fmoc-L-Lys(Mtt)-OH (3 eq, 328 mg, 2.1 mmol) with the coupling reagents (as described) were added. The resin was washed as before, Fmoc deprotected, and Fmoc-D-glutamic acid α-amide (3 eq, 223 mg, 2.1 mmol) for "Q" / "Gln" peptides *or* Fmoc-D-glutamic acid-OtBu-OH (3 eq, 223 mg, 2.1 mmol) for "E" / "Glu" peptides was added with the same coupling reagents as previously described. The Fmoc deprotection and coupling procedure was repeated as before using the same equivalencies with Fmoc-L-Alanine-OH. The Fmoc group of L-alanine was deprotected and coupled with 5(6)-carboxyfluorescein (2 eq, 132 mg, 1.4 mmol) for "FL" peptides or 5(6)-carboxytetramethylrhodamine (2 eq, 151 mg, 1.4 mmol) for "TAMRA" peptides, HBTU (2 eq, 133 mg, 1.4 mmol) and DIEA (4 eq, 0.122 mL, 2.8 mmol) in DMF (10 mL) shaking overnight. The resin was washed as before and the Mtt group of Lys was removed with 15 mL of a 1% TFA solution in DCM with shaking at ambient temperature for 20 min. This step was repeated until a total of 50 mL of 1% TFA was used. Fmoc-Gly-OH (3 eq, 156 mg, 2.1 mmol) was added to the resin with shaking at room temperature for 2 h. Some of the resin was split to yield 1 Gly on the Lys side chain. With the remaining resin, the Fmoc group was removed, and this coupling step was repeated 2X to yield a total of 3 Gly on the Lys side chain, or 4X to yield 5 Gly on the Lys side chain. The resin was washed as before and added to a solution of TFA/$H_2O$/TIPS (95%, 2.5%, 2.5%, 20 mL) with agitation for 2 h at ambient temperature. The resin was filtered and resulting solution concentrated *in vacuo*. The residue was trituated with cold diethyl ether.

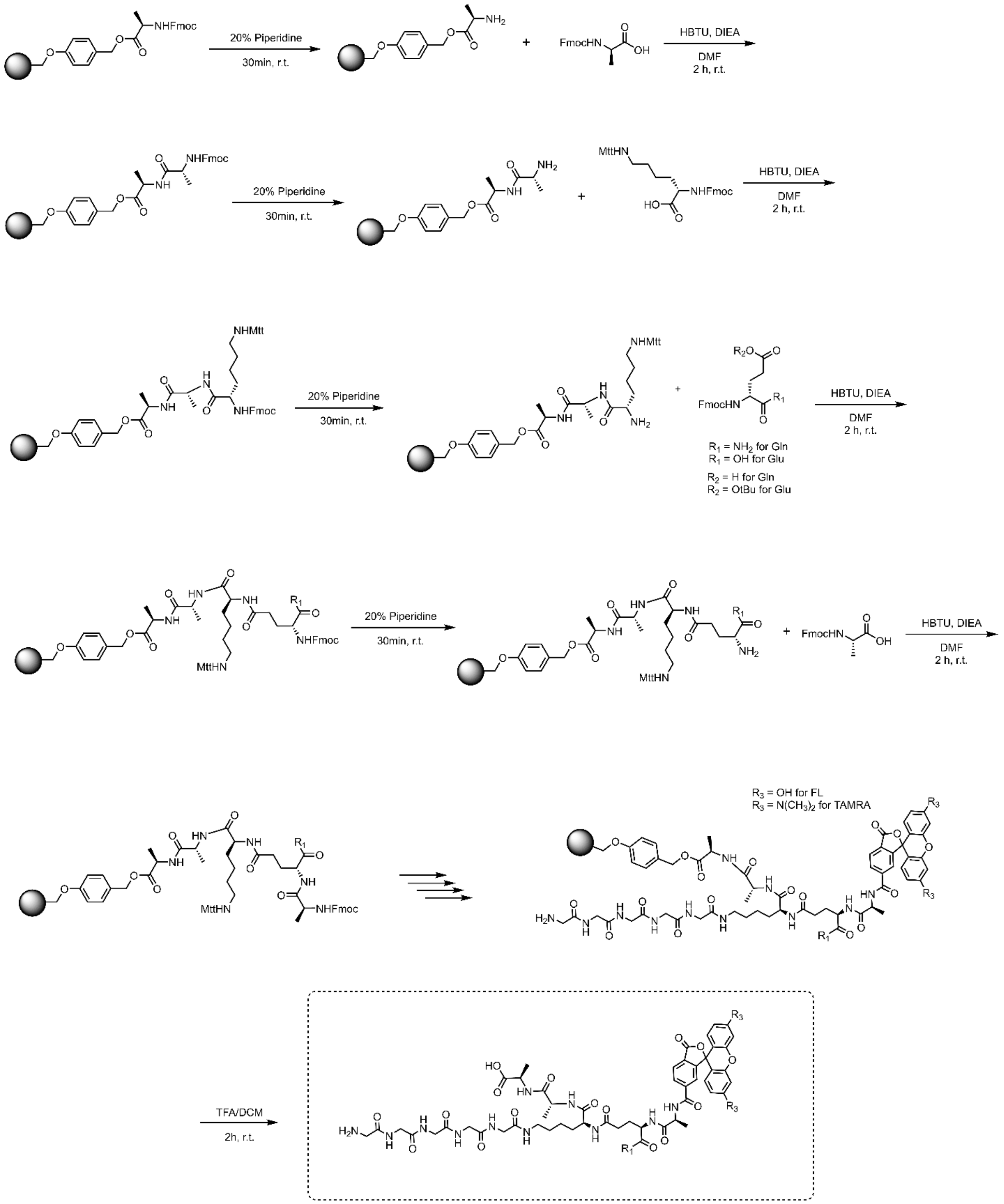
NHFmoc
20% Piperidine
30min, r.t.
$NH_2$
FmocHN
OH
HBTU, DIEA
DMF
2 h, r.t.
MttHN
NHFmoc
HO
NHMtt
$R_2O$
$R_1$
FmocHN
$R_1 = NH_2$ for Gln
$R_1$ = OH for Glu
$R_2$ = H for Gln
$R_2$ = OtBu for Glu
$R_3$ = OH for FL
$R_3 = N(CH_3)_2$ for TAMRA
$R_3$
$H_2N$
TFA/DCM
2h, r.t.

**Scheme S4. Synthesis of GGGK(TAMRA)**
Fmoc-Lys(Mtt)-Wang resin (250 mg) was added to a 25 mL peptide synthesis vessel with 20% piperidine in DMF for 30 min to remove the base-labile protecting group. The resin was washed with MeOH and DCM (3 x 15 mL each). Fmoc-Gly-OH (3 eq, 156 mg, 2.1 mmol), HBTU (3 eq, 200 mg, 2.1 mmol), and DIEA (6 eq, 0.183 mL, 4.2 mmol) in DMF (10 mL) were added to the reaction flask and agitated for 2 h at ambient temperature. The resin was washed as before and the Fmoc deprotection and coupling procedure was repeated as before using the same equivalencies for an additional 2 Gly. After washing, the Mtt group of Lys was removed with 15 mL of a 1% TFA solution in DCM with shaking at ambient temperature for 20 min. This step was repeated until a total of 50 mL of 1% TFA was used. 5(6)-carboxytetramethylrhodamine (2 eq, 151 mg, 1.4 mmol), HBTU (2 eq, 133 mg, 1.4 mmol) and DIEA (4 eq, 0.122 mL, 2.8 mmol) in DMF (10 mL) was added to the vessel with shaking overnight. The resin was washed as before and added to a solution of TFA/$H_2O$/TIPS (95%, 2.5%, 2.5%, 20 mL) with agitation for 2 h at ambient temperature. The resin was filtered and resulting solution concentrated *in vacuo*. The residue was trituated with cold diethyl ether.

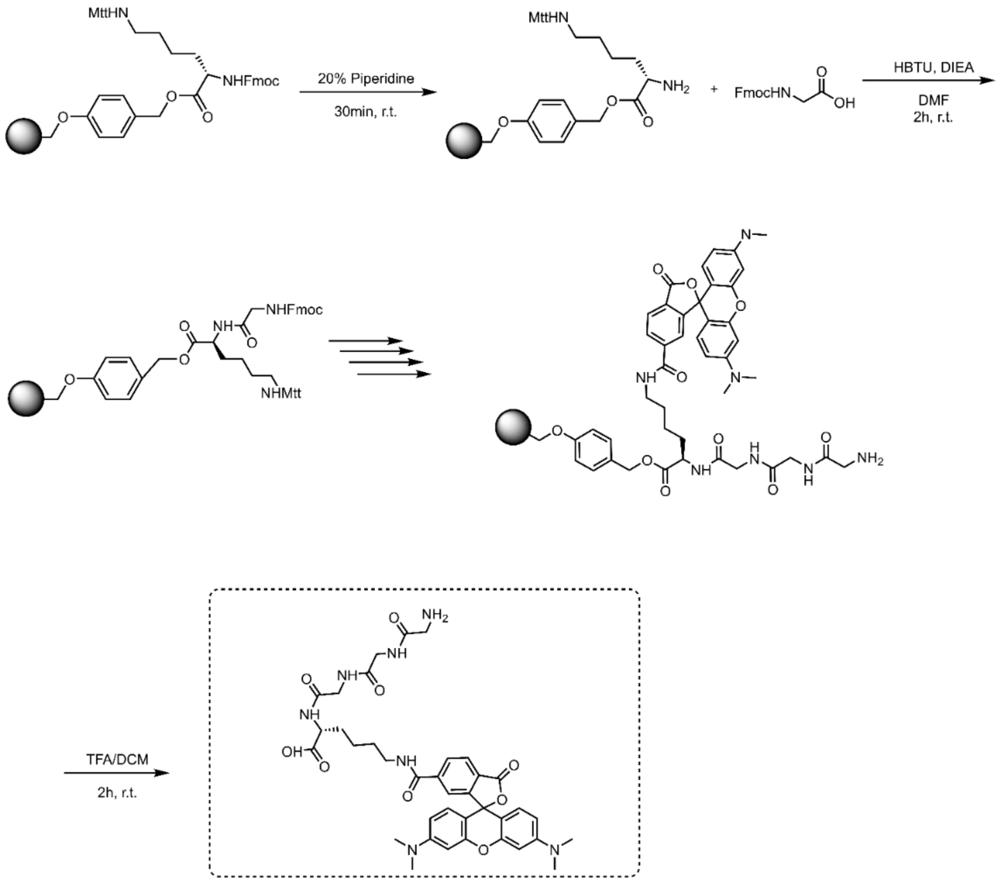

MttHN
NHFmoc
20% Piperidine
30min, r.t.
NH2
FmocHN
OH
HBTU, DIEA
DMF
2h, r.t.
NHMtt
TFA/DCM
2h, r.t.

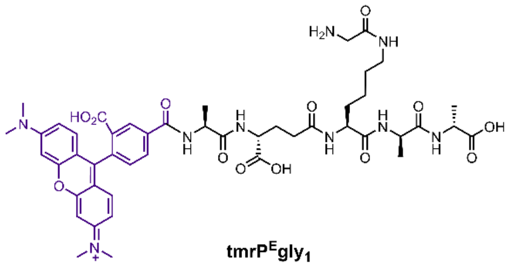

**tmrP$^{E}$gly$_1$**

### Mass Analysis:

Calculated M+2H+ = 479.7189
Found = 479.7186

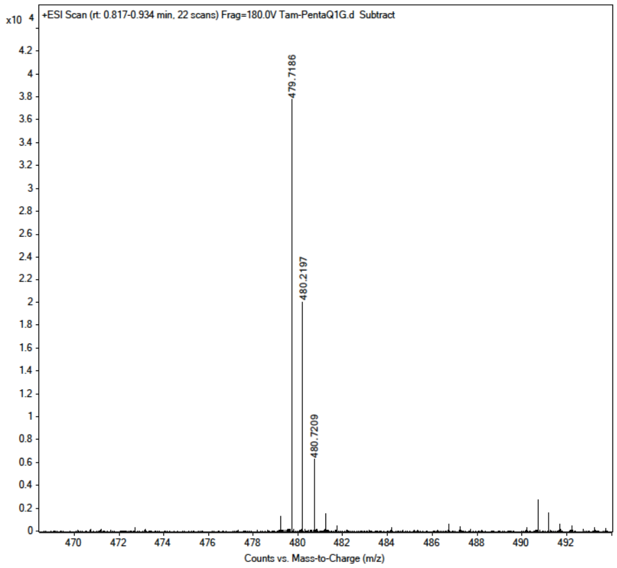


### RP-HPLC Analysis:

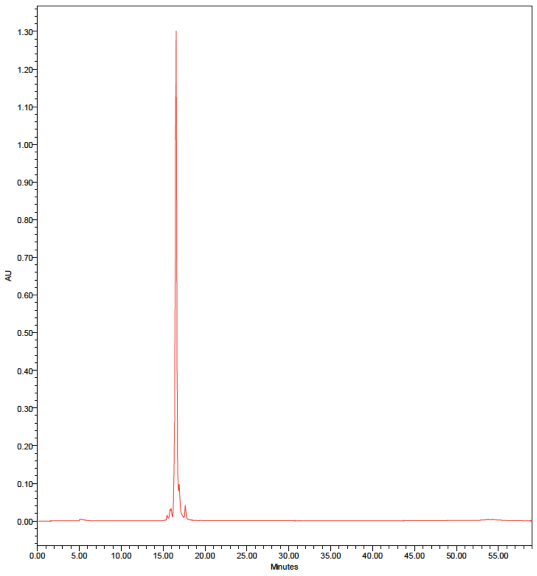
AU
Minutes

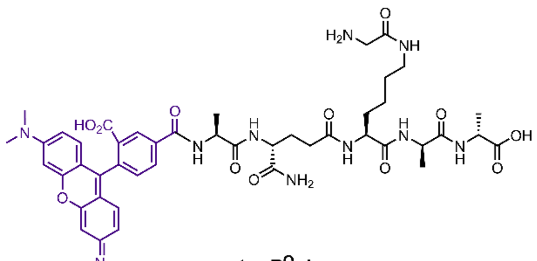


**tmrP$^{Q}$gly$_1$**

### Mass Analysis:

Calculated M+2H+ = 479.2269
Found = 479.2266

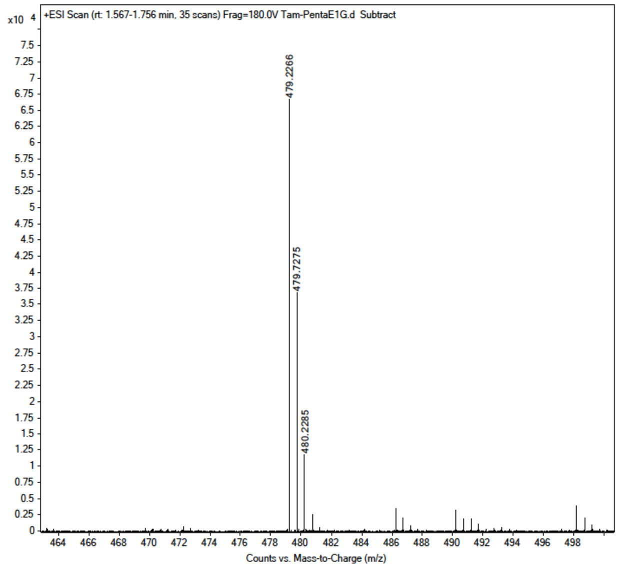


### RP-HPLC Analysis:

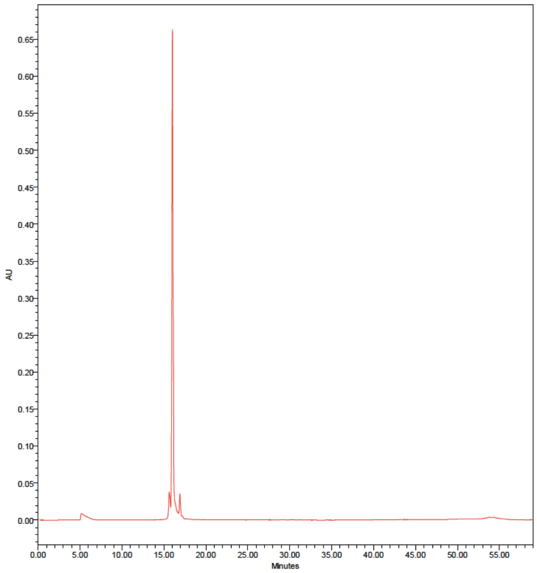
AU
Minutes
0.00
0.05
0.10
0.15
0.20
0.25
0.30
0.35
0.40
0.45
0.50
0.55
0.60
0.65
0.00
5.00
10.00
15.00
20.00
25.00
30.00
35.00
40.00
45.00
50.00
55.00

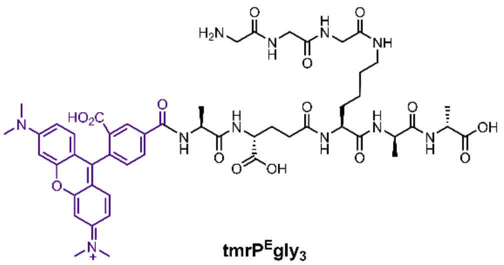

**tmrP$^{E}$gly$_3$**

#### Mass Analysis:

Calculated M+2H+ = 536.7404
Found = 536.7391

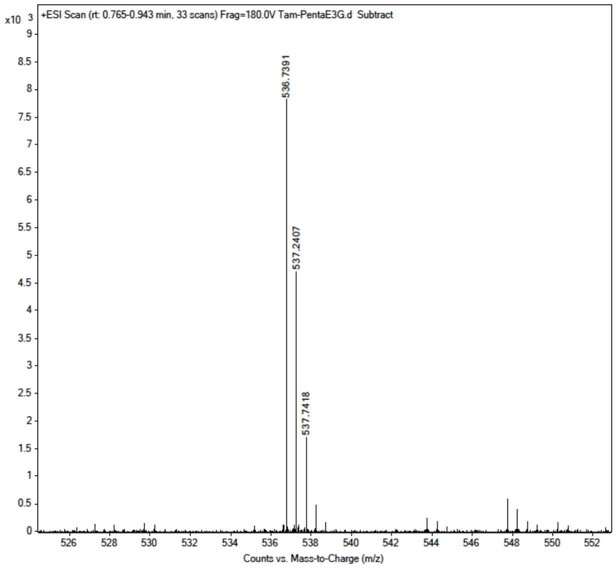


#### RP-HPLC Analysis:

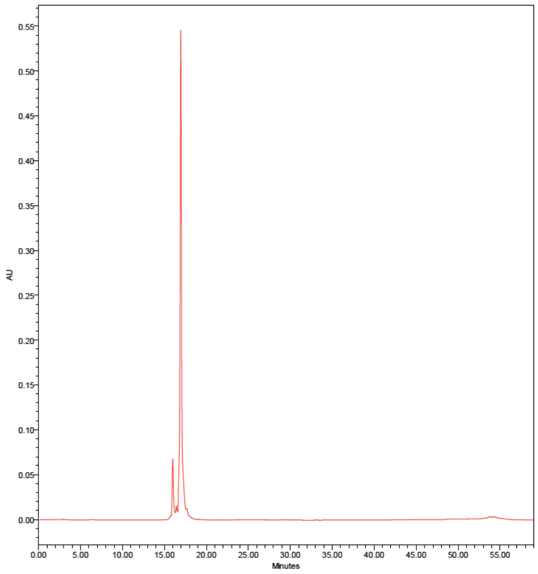
AU
Minutes
0.55
0.50
0.45
0.40
0.35
0.30
0.25
0.20
0.15
0.10
0.05
0.00
0.00
5.00
10.00
15.00
20.00
25.00
30.00
35.00
40.00
45.00
50.00
55.00

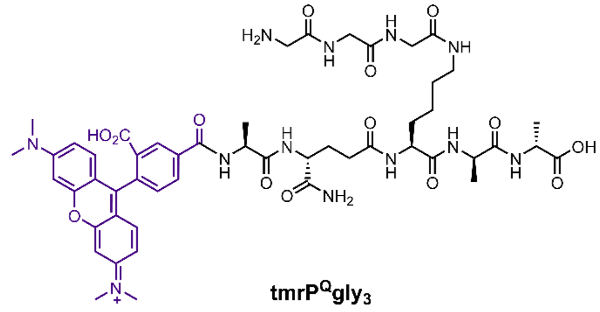


Mass Analysis:

Calculated M+2H+ = 536.2483
Found = 536.2478

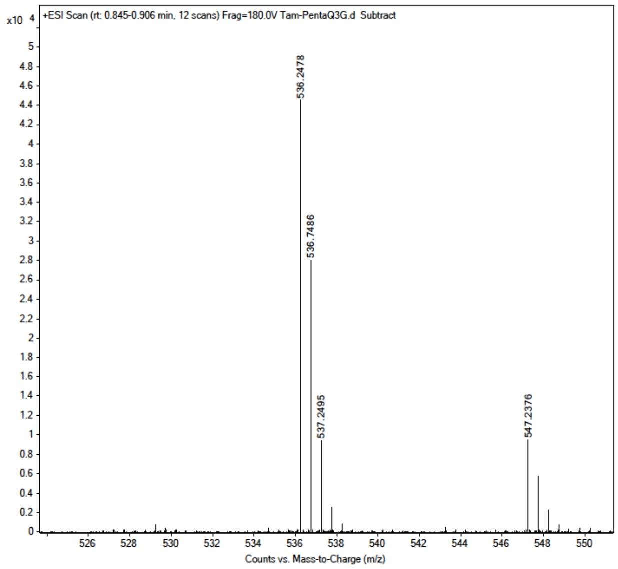


RP-HPLC Analysis:

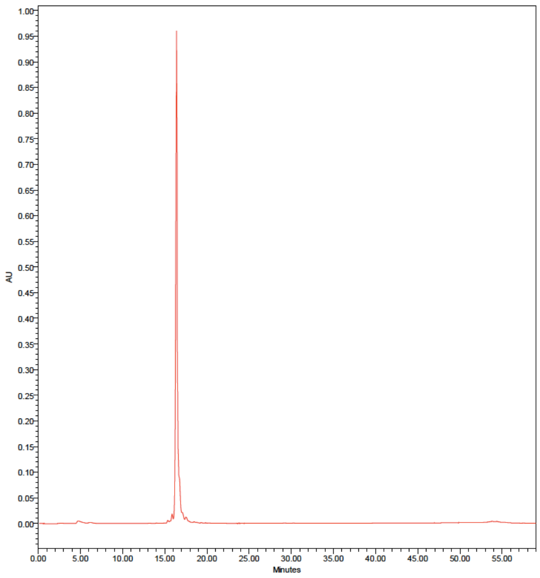

AU
Minutes

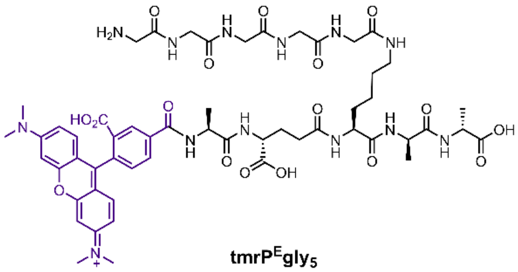


#### Mass Analysis:

Calculated M+2H+ = 522.7247
Found = 522.7239

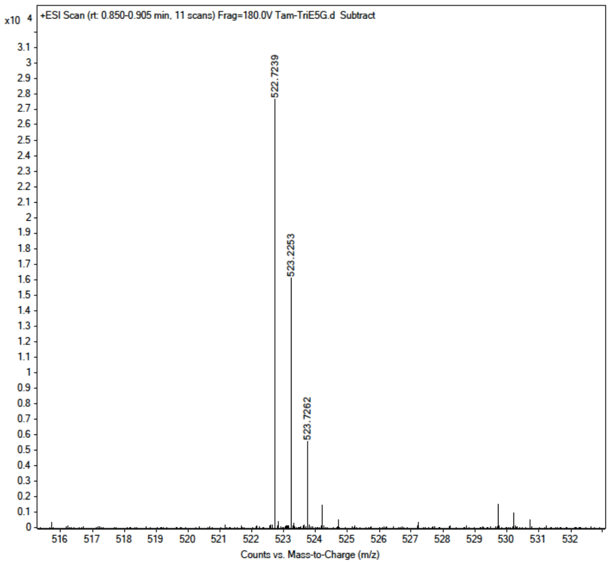


#### RP-HPLC Analysis:

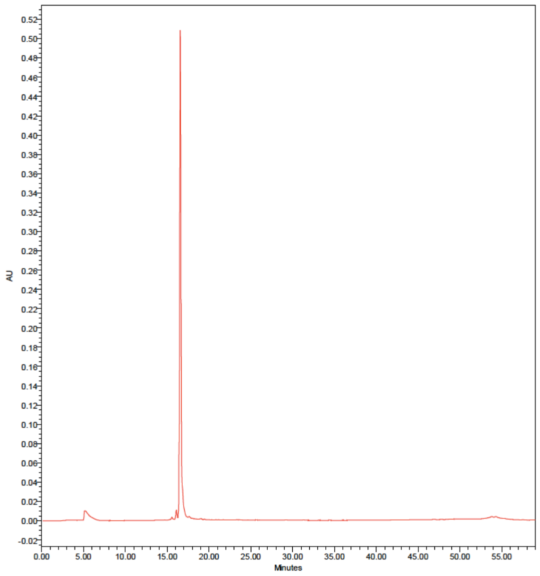
AU
Minutes
0.52
0.50
0.48
0.46
0.44
0.42
0.40
0.38
0.36
0.34
0.32
0.30
0.28
0.26
0.24
0.22
0.20
0.18
0.16
0.14
0.12
0.10
0.08
0.06
0.04
0.02
0.00
-0.02
0.00
5.00
10.00
15.00
20.00
25.00
30.00
35.00
40.00
45.00
50.00
55.00

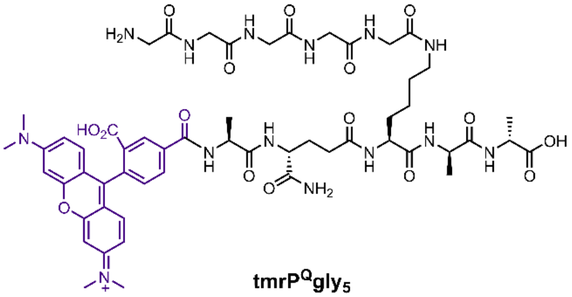

**tmrP$^{Q}$gly$_5$**

Mass Analysis:

Calculated M+2H+ = 522.2327
Found = 522.2314

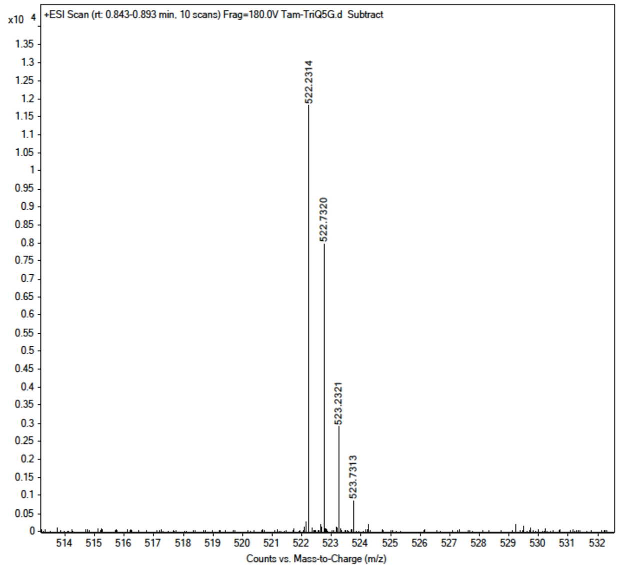

RP-HPLC Analysis:

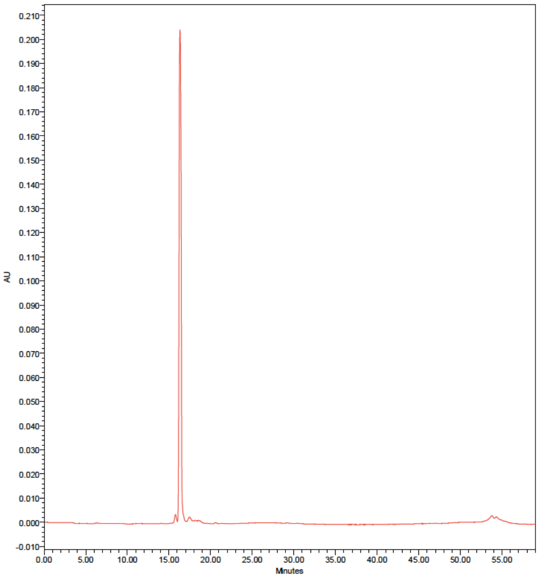

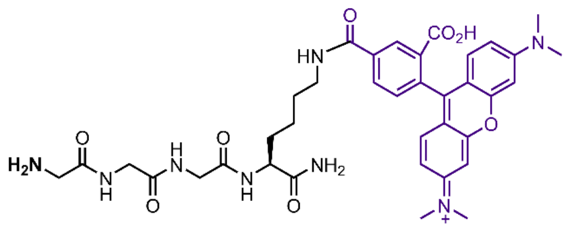

#### Mass Analysis:

Calculated M+H+ = 730.3200
Found = 730.3179

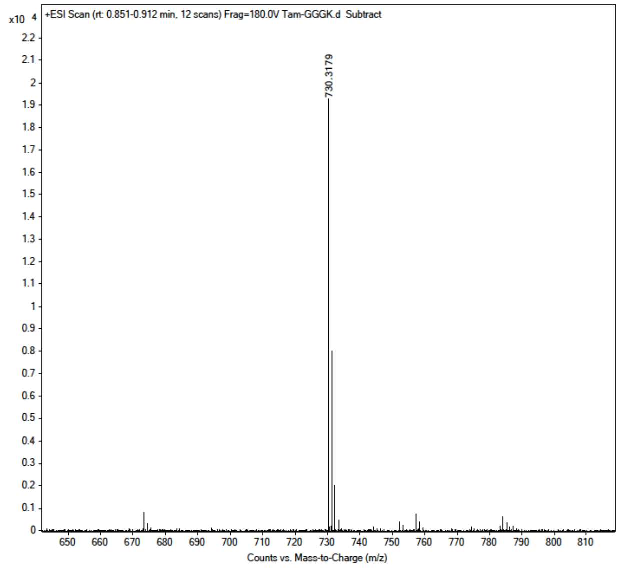


#### RP-HPLC Analysis:

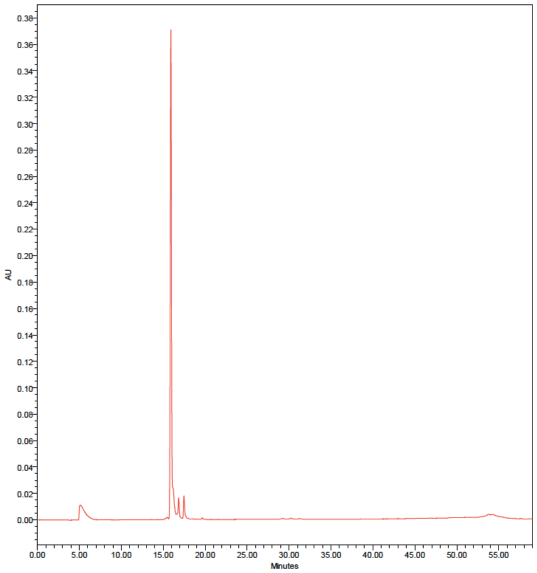

AU
Minutes
0.00
0.02
0.04
0.06
0.08
0.10
0.12
0.14
0.16
0.18
0.20
0.22
0.24
0.26
0.28
0.30
0.32
0.34
0.36
0.38
0.00
5.00
10.00
15.00
20.00
25.00
30.00
35.00
40.00
45.00
50.00
55.00

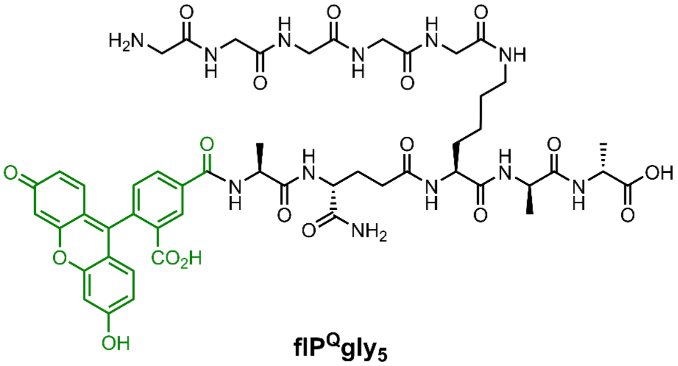

**flP$^{Q}$gly$_5$**

#### Mass Analysis:

Calculated M+H+ = 1132.4223
Found = 1132.4178

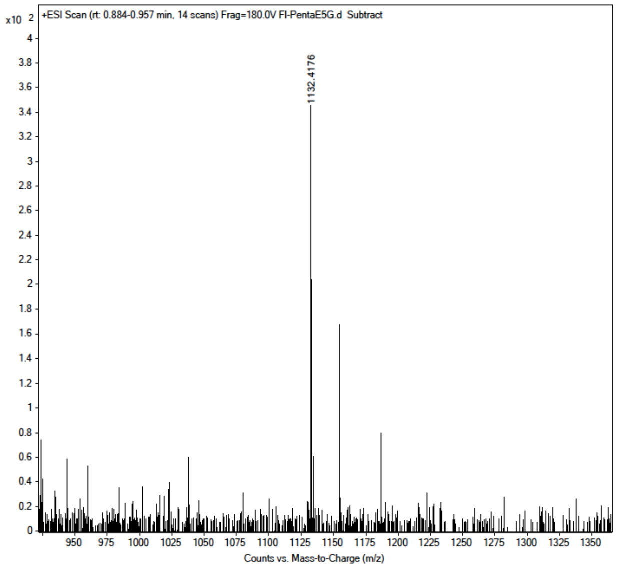


#### RP-HPLC Analysis:

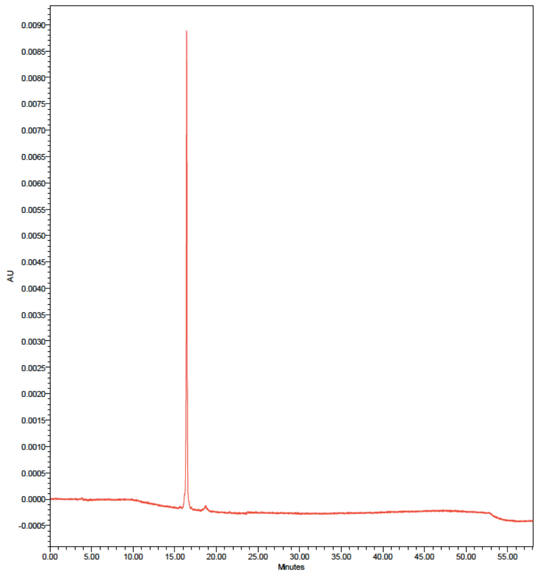

AU
Minutes

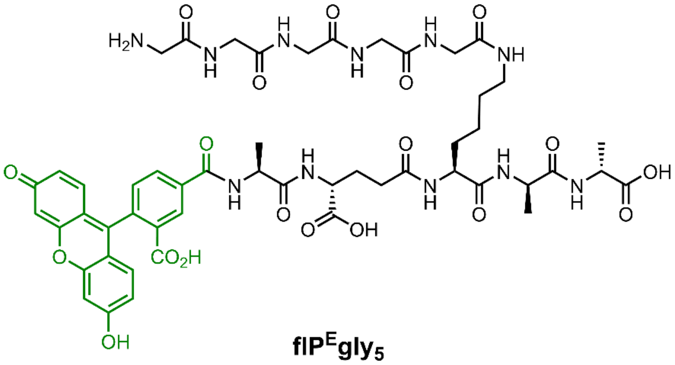

**flP$^{E}$gly$_5$**

Mass Analysis:

Calculated M+H+ = 1131.4383
Found = 1131.4329

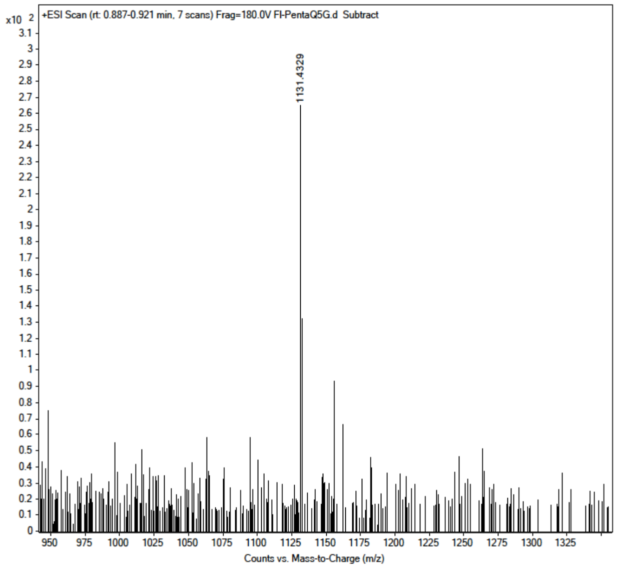

RP-HPLC Analysis:

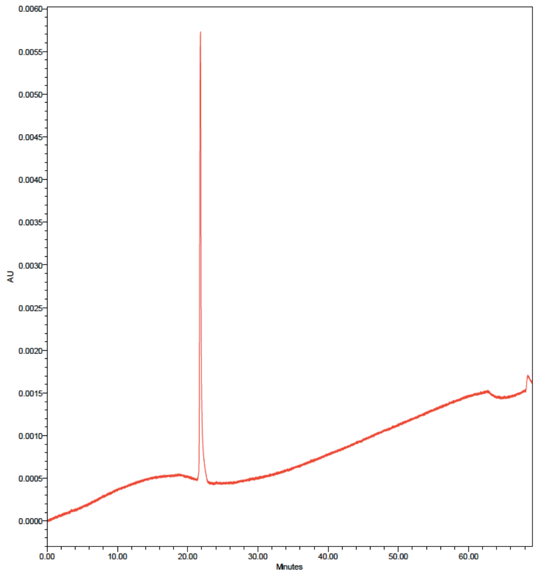

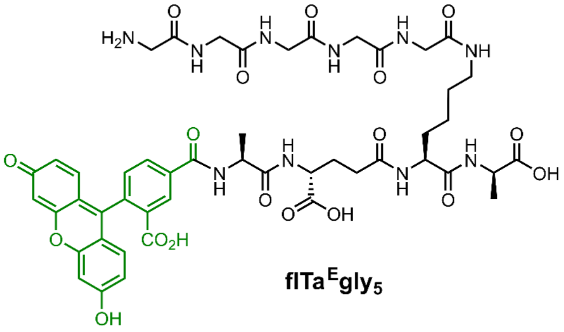

**flTa$^{E}$gly$_5$**

### Mass Analysis:

Calculated M+H+ = 1061.3852
Found = 1061.3718

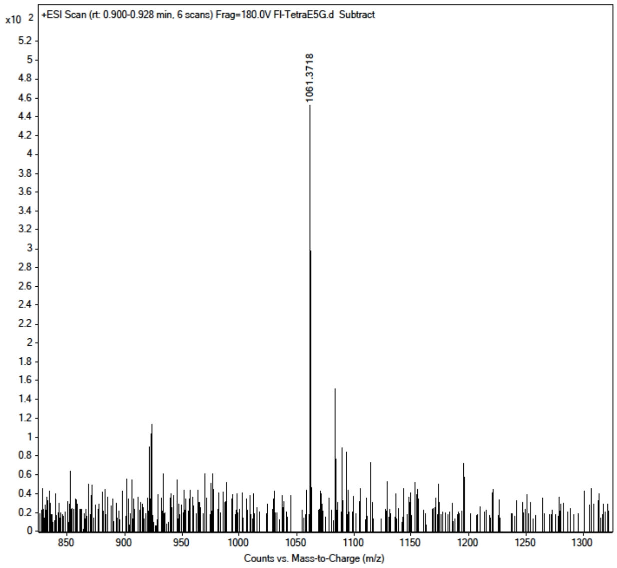

RP-HPLC Analysis:

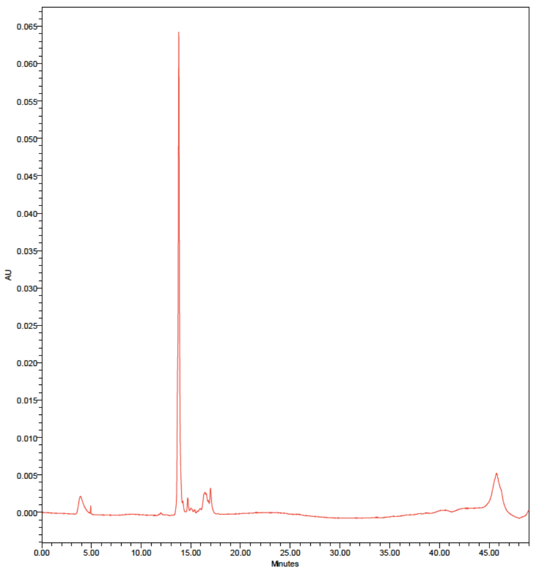

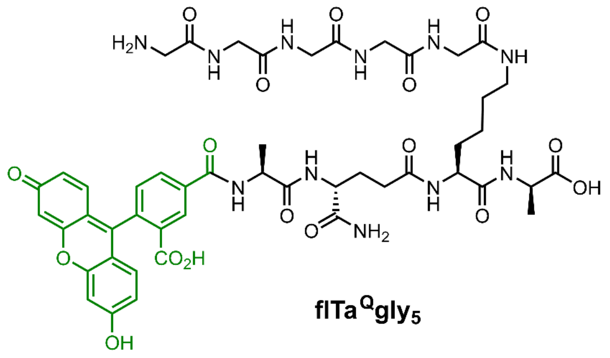


#### Mass Analysis:

Calculated M+H+ = 1060.4007
Found = 1060.3970

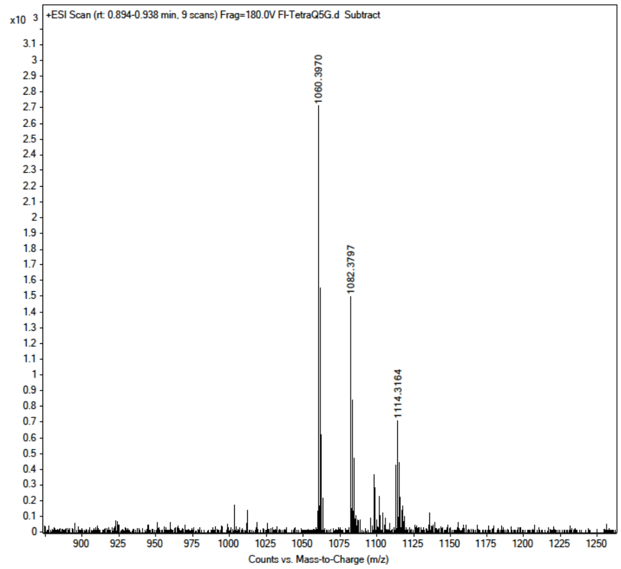

RP-HPLC Analysis:

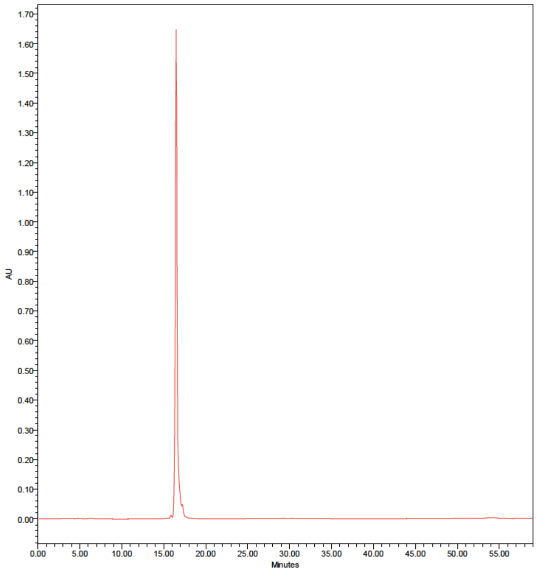

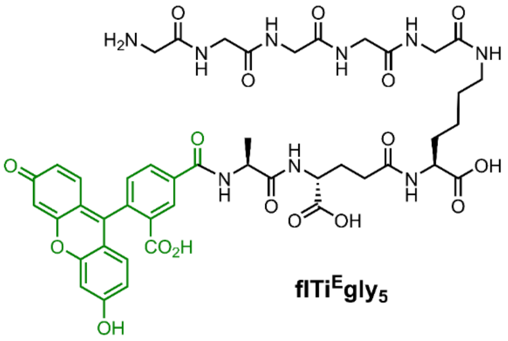

**flTi$^{E}$gly$_5$**

Mass Analysis:

Calculated M+H+ = 990.3481
Found = 990.3457

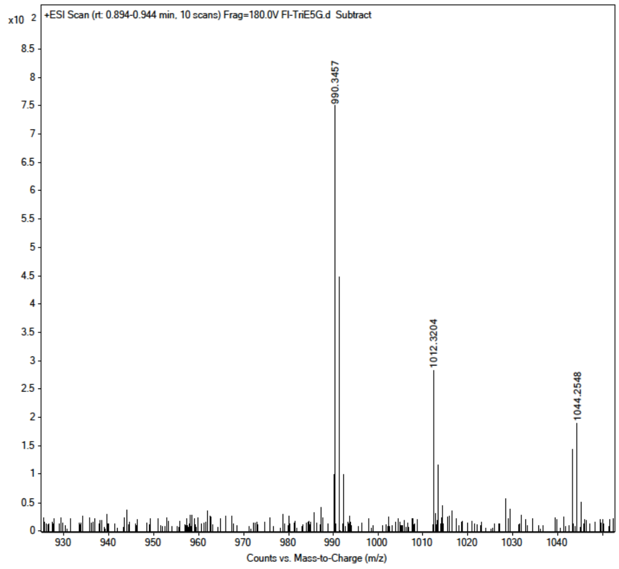

RP-HPLC Analysis:

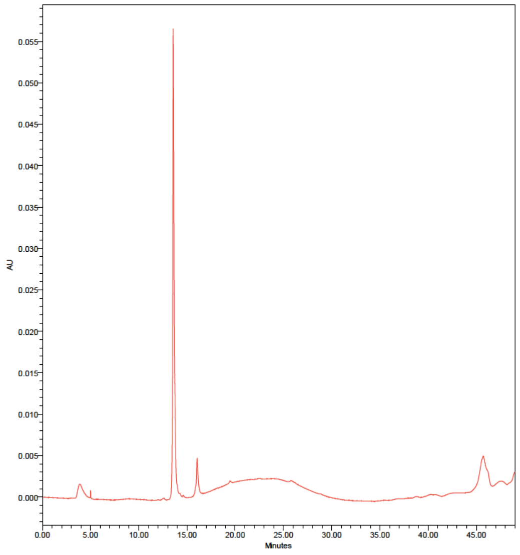

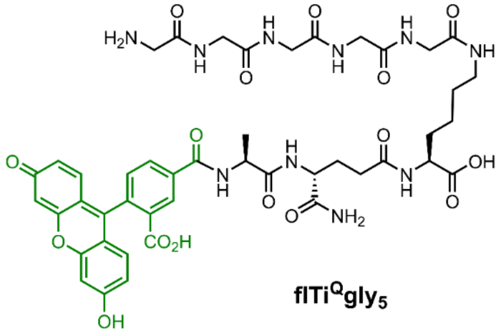

**flTi^Qgly$_5$**

### Mass Analysis:

Calculated M+H+ = 989.3641
Found = 989.3688

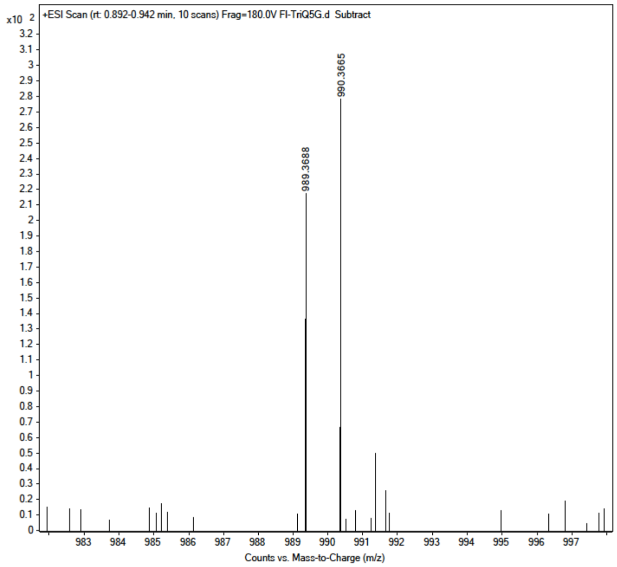

RP-HPLC Analysis:

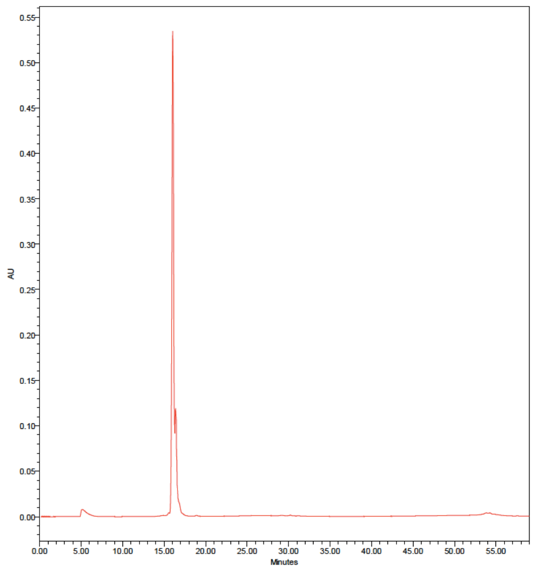
